# Metagenomic characterisation of fungal communities associated with Scots pine bark beetles: mechanisms of selective antagonism and monoterpene tolerance

**DOI:** 10.64898/2025.12.15.694374

**Authors:** Arunabha Khara, Sandipan Banerjee, Amrita Chakraborty, Jakub Dušek, Jiří Synek, Amit Roy

**Affiliations:** Czech University of Life Sciences Prague, Faculty of Forestry and Wood Sciences, Kamýcká 129, CZ – 165 21 Praha 6 – Suchdol, Czech Republic

**Keywords:** *Ips sexdentatus*, *Ips acuminatus*, fungal community, ITS2 amplicon sequencing, monoterpenes, scanning electron microscopy, enzyme assay, selective antagonism

## Abstract

Bark beetle-associated fungi contribute to beetle nutrition, detoxification, and interactions with conifer hosts; however, their composition and function across development and environments remain poorly understood. We characterised fungal communities of two pine-feeding *Ips* bark beetles, *I. sexdentatus* and *I. acuminatus*, in larval, pupal, and adult stages, and in wild versus laboratory populations, using high-throughput ITS2 amplicon sequencing combined with qPCR and functional assays. Both beetles harboured a stable core mycobiome dominated by *Kuraishia*, *Ogataea*, *Ophiostoma*, *Graphilbum* and *Cyberlindnera*, while adults showed species-specific differences and wild beetles, especially *I. acuminatus*, exhibited greater diversity than laboratory populations. Beetles shared more taxa with unfed control wood than with gallery wood, indicating acquisition during feeding and concurrent restructuring of the wood mycobiome. Monoterpene bioassays on beetle-associated yeasts revealed that mixtures of α-pinene, 3-carene and terpinolene suppressed growth more strongly than single compounds, suggesting synergistic inhibition. Yeasts selectively antagonised entomopathogenic fungi and expressed complementary cell-wall-lytic and digestive activities, consistent with combined roles in pathogen suppression and plant-polymer deconstruction. Our results show that *Ips* mycobiomes are conserved yet dynamic across life stages and environments, and emphasise the importance of multi-terpene and interaction assays for understanding bark beetle–fungus–conifer interactions.

**Importance:** Our study reveals that pine-feeding bark beetles co-evolved in close association with a stable core mycobiota that supports nutrient acquisition, detoxification, and chemical signalling, while additional fungal partners shift in response to beetle development, environment, and host context. By demonstrating higher mycobiome diversity in beetle larvae and wild populations, substantial overlap between beetle- and wood-associated fungi, differential sensitivity to monoterpene blends and selective antagonism of yeasts towards entomopathogenic fungi, our study unravels key ecological and mechanistic principles shaping beetle-fungus assemblages under conifers, paving the way for more in-depth functional investigations.

## Introduction

Bark beetles (Coleoptera: Curculionidae: Scolytinae) are among the most consequential forest pests, capable of triggering landscape-scale outbreaks and substantial economic losses (Singh et al., 2024). In Europe, climate change, combined with human-driven disturbances, has amplified the aggressiveness of pine-feeding species such as *Ips acuminatus* and *I. sexdentatus*, which now frequently colonise not only weakened but also seemingly healthy trees (Biedermann, 2019; Colombari, 2013; Faccoli, 2012; Pineau, 2017; Plewa, 2017). For instance, in the Czech Republic alone, bark beetle infestations on pinewood increased from ∼10,000 m³ in 2009 to nearly 80,000 m³ in 2019 (Liška, 2021). Climate change has exacerbated bark beetle outbreaks by weakening tree defences through rising temperatures, altered precipitation, and frequent drought and heat events (Marini et al., 2017; Raffa et al., 2008). Drought in particular reduces host tree vigour, enabling beetle populations to surpass epidemic thresholds (McNichol et al., 2022).

Successful colonisation of pine hosts requires beetles to overcome robust constitutive and induced defences. Physical barriers (lignin- and suberin-rich tissues) and chemical defences (phenolics and terpenoids) impede invasion and development (Mumm & Hilker, 2006). Bark beetle-fungus mutualism is one such crucial interaction in which the fungal partners have evolved specific adaptations that enable beetles to thrive in the toxin-laden conifer environment and utilise the nutrient-deficient phloem (Linnakoski, 2012). Microbial communities residing inside (i.e., gut, hemolymph, or any other specialised tissue) produce vital enzymes that have a specialised function in the insect, including bark beetles (Blow, 2019). For instance, in ambrosia beetles, the fungal symbionts reside in specialised structures (i.e., mycangia), whereas in *Ips* beetles, fungi produce spores that can easily attach to beetle bodies (Francke-Grosmann, 1967; Six, 2011). These spores contain a well-protected sheath that prevents digestion inside the beetle gut (Seifert, 2013). These fungal associates contribute to beetle fitness through multiple, complementary functions, including nutrient acquisition, detoxification of plant defence compounds, and the production of volatile compounds to facilitate beetle communication (Douglas, 2015). For example, the fungal partners degrade the complex biopolymers of wood, such as hemicellulose, lignin, and cellulose, making them nutritionally accessible to their host beetles (Kirk, 1984; Valiev, 2009). Several fungal associates of the Eurasian spruce bark beetle (*Ceratocystis polonica*, *Grosmannia europhioides*, *Grosmannia penicillate*) facilitate its attack on healthy trees through their involvement in the degradation of spruce phenolic compounds (flavonoids, stilbenoids) (Hammerbacher, 2013; Zhao, 2019). Subsequently, the predominant fungal isolates, such as *Grosmannia*, *Ophiostoma*, and *Endoconidiophora* of bark beetles, can utilise tree monoterpenes as carbon sources to generate aggregation pheromones, facilitating mass beetle attacks (Kandasamy, 2023). Similarly, fungal isolates from bark beetles convert such aggregation volatiles (trans-verbenol and cis-verbenol) into non-aggregation volatiles, thereby minimising intraspecific competition (Hunt, 1990; Netherer, 2021). Fungal mutualists also provide bark beetles with protection against other pathogens. For example, the symbiotic fungus *Leptographium abietinum*, which is associated with the spruce beetle, can inhibit the entomopathogenic fungus *Beauveria bassiana* (Davis, 2019). Therefore, it is evident that fungal mutualists play a crucial role in the survival of bark beetles under challenging environments (Liu, 2022).

Despite extensive work on symbioses in several model bark beetles, including the red turpentine beetle, mountain pine beetle (Khadempour, 2012; Taerum, 2013) and Eurasian spruce bark beetle (Chakraborty, 2023; Veselská, 2023), comparatively little is known about the mycobiomes of the pine-feeding *Ips* species *I. acuminatus* and *I. sexdentatus*, particularly across ontogeny. Furthermore, the understanding of the fungal communities across potential host–environment filters, such as laboratory versus wild conditions, and the surrounding wood mycobiome, remains insufficiently studied for these taxa. Addressing these gaps is critical, based on emerging evidence that beetles can both acquire microbes from, and reshape, the fungal communities of their gallery environment.

Hence, in our study, we identified the fungal communities associated with *I. acuminatus* and *I. sexdentatus* across different life stages using high-throughput fungal ITS2 amplicon sequencing. We compared lab-bred and wild-collected adults to evaluate environmental contributions to fungal assemblages. We further tested the responses of a few selected culturable beetle-associated yeasts to ecologically relevant monoterpene exposures and a mixture that partially mimics host tissue defence chemistry. We hypothesise (1) both *Ips* species harbour a conserved core mycobiome across development, overlaid by life-stage-specific transient taxa; (2) environmental context (laboratory vs. wild) and the host-tree/wood microbiome structure beetle-associated fungal assemblages, yielding higher diversity in wild beetles; (3) monoterpene exposures might inhibit or delay fungal growth, with mixtures might exerts more potent effects than single compounds, reflecting synergistic toxicity and thus acting as ecological filters that favour more tolerant, functionally important symbionts, (4) fungal symbionts themselves restrict and shape the beetle mycobiome through competitive and facilitative interactions. By integrating community profiling followed by targeted interaction and enzymatic assays of selected fungal symbionts, we link mycobiome structure to putative ecological functions in nutrition, detoxification, and chemical signalling, and generate a set of testable hypotheses for future evolutionary studies on *Ips* beetles.

## Materials and Methods

### Bark beetle rearing and collection

Multiple logs (dbh ∼40 cm) infested with two adult *Ips* pine beetles (*Ips sexdentatus* and *Ips acuminatus*) (Coleoptera: Curculionidae: Scolytinae) were collected from different pine trees at the Rouchovany forest area (49.0704° N, 16.1076° E) in the Czech Republic, 2020. Published literature was utilised for the taxonomic identification of both adult beetles. The infested logs collected from the forest were immediately brought to the debarking room, and the adult beetles were collected using surface-sterilised tweezers. The collected adult beetles were then transferred to fresh uninfested logs collected from the same forest area and reared up to the F_2_ generation under laboratory conditions (day temperature, 25°C; night temperature, 19°C; relative humidity, 60%) as described in our previous studies (Khara et al., 2024). Approximately 50 samples, representing each life stage (larvae, pupae, and adults) from several infested logs (at least 10 cm below the edges to minimise contamination), were collected in 50 ml conical tubes. These conical tubes were snap-frozen with liquid nitrogen and kept at −80℃ for future metagenomic sample preparation. The gallery wood (comprising both mother and larval galleries) was also collected from the same F_2_ generation infested logs, following a standard protocol (Hulcr, 2022). Similarly, uninfested, fresh wood samples (unfed phloem tissue) were collected as a control for the wood. Both wood samples were collected using surface-sterilised blades, which were kept in conical tubes with RNAlater solution at −80℃. The size of the individual wood sample replicate was 0.5 by 1 cm sections. In the following year, 2021, multiple infested logs from different pine trees were collected from the same Rouchovany forest area. The adult beetles and their respective gallery wood samples were collected from the same infested logs as mentioned above.

### DNA extraction

All the bark-beetle samples (larvae, pupae, adults) were randomly selected, and the disinfection of the beetles was carried out using 70% ethanol for 1 min, followed by washing in sterile water. This process was repeated three times to remove any surface contaminants. Five biological replicates from each developmental stage and four from each wood sample were selected for fungal ITS2 amplicon sequencing at Novogene, China. Because of variation in size, the individuals in each biological replicate differed for both species; for *I. sexdentatus,* one individual per replicate was used. However, for *I. acuminatus* larvae (4 larvae/replicate), pupae (2 pupae/replicate), and adults (5 adults/replicate) were used. The DNA extraction and quantification have been previously reported in our studies (Khara et al., 2024). Briefly, the samples were subjected to total DNA extraction using the Macherey-Nagel NucleoSpin Soil DNA kit. DNA extraction from wood samples (100-120 mg per replicate) was performed using the DNeasy Plant Mini Kit (Qiagen, Germany). The extracted DNA samples underwent purification to eliminate any potential PCR inhibitor using the DNeasy Powerclean Pro cleanup kit (Qiagen, Germany). The purified DNA samples were quantitatively and qualitatively assessed by 1% agarose gel electrophoresis and Qubit 2.0 Fluorometer (Thermo Scientific).

### Amplicon sequencing

Diluted purified DNA (1ng/µl), fungal ITS2 region primers (ITS3, ITS4) (White, 1990) tagged with unique barcodes, Phusion High-Fidelity PCR Master Mix (New England Biolabs) were used to initiate PCR reactions. Subsequently, a no-template PCR reaction was performed as a negative control to verify the absence of contamination. The PCR amplicons were visualised on a 2% agarose gel, and the Qiagen Gel Extraction Kit (Germany) was used to purify the amplified products. Sequencing libraries were constructed with equidensities of amplified products using NEBNext Ultra II DNA Library Pre-Kit (Illumina). Qubit 2.0 Fluorometer (Thermo Fisher Scientific) and Agilent Bioanalyser 2100 system were used to check the quality and quantity of the developed libraries. Finally, 250 bp paired-end reads were generated from the libraries using the Illumina NovaSeq 6000 sequencing platform.

### Bioinformatic Data Analysis

#### Data processing and species annotation

Bioinformatic analysis of the mycobiome was executed using QIIME2 (version 2022.2) (Bolyen, 2019). After the removal of the primer and barcode sequences, Illumina pair-end raw reads were generated using FLASH (V1.2.11) (Magoč, 2011). The quality of the assembled raw reads was checked using the ‘fastp’ software to assemble high-quality clean reads. Chimeric sequence detection and removal were executed using VSEARCH software (Rognes, 2016) to facilitate downstream bioinformatic analyses. The DADA2 module in QIIME2 (Callahan, 2016) was used to remove sequences with fewer than five reads to generate ASVs (Li, 2020) and the ASV abundance table. UNITE database (Nilsson, 2019) and Classify-sklearn module (Bokulich, 2018) of QIIME2 (version 2022.2) (Bolyen, 2019) were used for annotating taxa to the ASVs. The core fungal ASVs that were present in at least 60% of each beetle sample group were taken into consideration to avoid the inclusion of any transient or accidentally occurring species during sample collection and processing.

#### Alpha diversity

Alpha diversity indices were used to assess the richness and diversity of the fungal community within a sample. The estimation, representation of indices like community richness (Chao1) (Magurran, 2013), evenness (Pielou) (Magurran, 2013) and diversity (Shannon, Simpson) (Magurran, 1988), Good’s coverage (sequence depth) (Chao, 1988) were done using QIIME2 (version 2022.2) and R software (Team, 2019). The statistical analysis between each group was performed using the Kruskal-Wallis pairwise group test.

#### Beta diversity

The variation in fungal diversity between different samples was estimated using the UniFrac distance metric (Lozupone, 2011) with QIIME2 (version 2022.2). The R software package was used to represent non-metric multidimensional scaling (NMDS) based on UniFrac distance measurement (Oksanen, 2015). The determination of significant differences in the mycobiome was performed using ADONIS and ANOSIM functions (Anderson, 2001; Clarke, 1993) in QIIME2 (version 2022.2). The ADONIS analysis is a non-parametric multivariate variance test used to reveal significant differences among sample groups (Stat, 2013). Subsequently, an ANOSIM function analysis determines whether the variation between different groups is greater than the within-sample group variation (Chapman, 1999). The differentially abundant fungal communities in the various sample groups were identified using Metastats analysis (employing the false discovery rate-FDR) (Paulson, 2011) and a t-test (p-value < 0.05) (D’Argenio, 2014). Statistically significant biomarkers were identified between the tested samples using LEfSe (linear discriminant analysis effect size) analysis (Segata, 2011). The probable functional/ecologically categorised profile was obtained using FUNGuild software (Nguyen, 2016).

### Quantitative PCR assay

The relative abundance of selected fungal taxa was measured using a quantitative PCR (qPCR) assay, following the optimised protocol (Khara, 2024). For *I. sexdentatus* samples, six individuals were used per replicate for each developmental stage (larvae, pupae, and adults). For *I. acuminatus*, only adults were analysed due to limited sample availability. Four biological replicates were prepared for each group. For qPCR assay, one universal fungal ITS2 primer and four fungal genus-specific primers were used (Supplementary Table 1). Fungal genus-specific primers for *Nakazawaea, Kuraishia, Ophiostoma*, and *Ogatea* were designed in-house, using available ITS gene sequences from NCBI. The primer specificity is determined by sequencing the amplified products and confirming the identity through BLAST analysis. qPCR reactions were conducted in 10 μL volumes, which included 4 μL of template DNA (10 ng μL⁻¹), 5 μL of SYBR® Green PCR Master Mix (Applied Biosystems), and 0.5 μL of each forward and reverse primer (10 μM). Amplification was performed on a QuantStudio thermocycler with the following conditions: an initial denaturation at 95 °C for 5 minutes, followed by 40 cycles of denaturation at 95 °C for 15 seconds and annealing/extension at 60 °C for 30 seconds. For normalisation, we used housekeeping genes that showed stable expression across samples: β-tubulin for *I. sexdenatatus* samples (Sellamuthu, 2021) and elongation factor 1-α (*EF1α*), ribosomal protein (*RPL7*) for *I. acuminatus* samples (unpublished data). We calculated relative quantification (RQ) using the 2⁻ΔΔCt method, where ΔCt indicates the difference between the Ct value of the target taxon and the reference gene. This method estimates fold changes in target abundance compared to the reference gene, avoiding inconsistencies that can occur when using the total fungal population as a denominator (Navidshad, 2012). We conducted statistical analysis of relative abundance in R (version 4.3.1) using ANOVA, followed by post-hoc tests and t-tests to identify significant differences between groups (Team, 2019). Furthermore, we correlated the qPCR results with the abundance data from metagenomic sequencing.

### Fungal cultures and identification

Fungal species were isolated from healthy pine beetles as well as pine wood samples. The beetles were disinfected using 70% ethanol for 1 minute, followed by three washes in sterile water to remove any surface contaminants. The beetles were then crushed in PBS solution, and serial dilutions (10^-1^ to 10^-6^) were prepared. The dilutions were spread on Potato Dextrose Agar (PDA) plates and incubated at 28 °C for 5 days. The wood samples were cut into small sections of 0.5cm and placed on the PDA plates. The fungal growth observed was subcultured thrice to obtain pure culture isolates. Similarly, the pathogenic fungal mycelia from the surface of the dead beetles were extracted and cultured on PDA plates. The identification of the fungal isolates was performed using Sanger sequencing, and the identity was analysed by NCBI BLAST. The sequences were submitted to the NCBI GenBank database with accession numbers PX491830-PX491839 (Supplementary Table 2)

### Monoterpene Bioassay

Four yeast strains (*Yamadazyma mexicana*, *Nakazawaea holstii*, *Kuraishia molischiana*, and *Cyberlindera mississippiensis*) isolated from two pine-feeding beetles were used in the bioassay with monoterpenes to understand their impact on yeasts. We assayed three selected monoterpenes (α-pinene, 3-carene, and terpinolene) and a monoterpene blend (a mixture of α-pinene, 3-carene, and terpinolene) at varying concentrations to mimic the chemical exposure conditions in nature within the host environment. Monoterpene concentrations were considered and calculated in accordance with earlier publications (Musso et al., 2023). For α-pinene, we used concentrations of 100 ng µL⁻¹, 500 ng µL⁻¹, and 1500 ng µL⁻¹; for 3-carene, 100 ng µL⁻¹, 500 ng µL⁻¹, and 1000 ng µL⁻¹; and for terpinolene, 50 ng µL⁻¹, 100 ng µL⁻¹, and 200 ng µL⁻¹. The monoterpene blend consisted of 1500 ng µL⁻¹ α-pinene, 1000 ng µL⁻¹ 3-carene, and 100 ng µL⁻¹ terpinolene. Dimethyl sulfoxide: DMSO 0.5% (v/v) was utilised as the control solvent since a concentration of that amount was required to assist in dissolving the monoterpenes in potato dextrose broth (PDB) medium. Overnight yeast cultures adjusted to an optical density (OD₆₀₀) of 0.5 were used to inoculate for the growth assay. Spectrophotometric measurements at OD₆₀₀ were taken at fixed intervals of time. The experiments were carried out in triplicate to facilitate reproducibility and consistency of data.

### Scanning electron microscopy (SEM)

#### Gut colonisation of fungi

To explore *I*. *sexdentatus* gut microbial colonisation, the dissected midgut and hindgut were fixed in 2.5 % glutaraldehyde in 0.2 M cacodylate buffer (pH 7.0) at 4°C for 7 days. The gut samples were then dehydrated using a varying concentration of ethanol wash (35%, 50%, 70% and 96%) with an incubation period of 10 minutes in each step. The samples were fixed to a cylinder with electron-conductive double-sided adhesive carbon tabs (EM-Tec CT6, Micro to Nano) and coated with gold (10 nm thickness) using a JFC 1300 Auto Fine Coater (JEOL). The samples were then observed under a JSM-IT500HR InTouchScope™ scanning electron microscope (JEOL) at an accelerating voltage of 3 kV with a working distance of 11-13.1 mm.

#### Biofilm formation

The biofilm-forming capability of the gut isolates was also examined under SEM. The four yeast isolates were grown in Potato Dextrose broth (PDB), and also starch, chitin, gelatin, cellulose, laminarin, pectin, and xylan supplemented broth (Supplementary Table 3) in static condition for 24-48 h at 28°C where small pieces of glass chips (1 × 1 cm) were placed at the bottom of the conical flask. After incubation, the yeast film formed on the glass chips at the lower portion of the media was collected and gently washed twice with sterile PBS (1X), followed by dehydration with ethanol gradients (5-100% at 5% increments for 1 min each). The samples were fixed to a cylinder, coated, and examined under SEM using the same parameters as previously mentioned.

#### Antifungal activity

To decipher the antifungal activity of the yeast isolates against the entomopathogenic fungal (EPF) strains and non-entomopathogenic fungal (NEPF) strains (Supplementary Table 2), mycelial morphologies were visualised under SEM. Fungal mycelia of pathogenic and non-pathogenic strains, as well as yeast symbionts, were obtained from pure cultures grown on PDB for 7 days and overnight (Optical density, OD, 0.5 at 600 nm), respectively. Alternatively, co-cultures of EPF-yeast and NEPF-yeast were prepared by inoculating EPF or NEPF grown on PDB for 7 days at 28°C with an overnight-grown yeast culture (optical density, OD, 0.5 at 600 nm) in PDB and incubating for 48 h at 28°C. The experimental setup included one EPF or NEPF that was co-cultured with one yeast at a time. The samples were fixed overnight in 2.5% glutaraldehyde in 0.2 M cacodylate buffer (pH 7.0) at 4°C. Subsequently, fixed mycelia were dehydrated using a series of ethanol washes with increasing concentrations of ethanol (10-95%) with a five-minute incubation for each step.

#### Enzyme production assay

The ability of the yeast isolates to produce antifungal enzymes (chitinase, protease, and β-glucanase) and digestive enzymes (amylase, cellulase, pectinase, and xylanase) was evaluated based on standard protocols (Danso et al., 2022; Dhayalan et al., 2022; Jacobs & Gerstein, 1960; Kuddus & Ahmad, 2013; Muriithi et al., 2022; Teather & Wood, 1982; Wu et al., 2018). Four yeast isolates were grown on specific agar media (Supplementary Table 3) and incubated at 28°C for 48 hours. The antifungal and digestive enzyme-producing capabilities of the gut isolates were assessed using a qualitative index (QI).

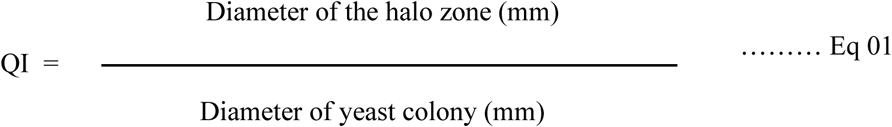

#### Estimation of antifungal enzymes

Selected antifungal enzymes (chitinase, protease, and β-glucanase activities) produced by the yeast isolates were quantified (U mL⁻¹) using standard substrate assays. Chitinase activity was assayed following (Song et al., 2017) with 1% (w/v) colloidal chitin in phosphate buffer (pH 7.0) as the substrate; one unit (U) was defined as the amount of enzyme releasing 1 mM N-acetylglucosamine (GlcNAc) per hour at 28 °C. Protease activity was determined using 1% (w/v) casein in phosphate buffer (pH 7.0), following published protocol (Cupp-Enyard, 2008), where one unit was defined as the amount of enzyme liberating 1 µM tyrosine equivalents from casein at 28 °C. While β-glucanase activity was measured according to published protocol (Ghose, 1987), using 1% (w/v) laminarin in phosphate buffer (pH 7.0) as the substrate, one unit was defined as the amount of enzyme releasing 1 mM glucose per minute at 28 °C. For each assay, the corresponding reaction mixture prepared with uninoculated production medium served as the control.

#### Estimation of digestive enzymes

Selected digestive enzymes (amylase, cellulase, pectinase, and xylanase activities) were quantified (U mL⁻¹) by the dinitrosalicylic acid (DNS) assay. Amylase was assayed according to (Miller, 1959) using 1% (w/v) soluble starch in phosphate buffer (pH 7.0) as the substrate; one unit (U) was defined as the amount of enzyme releasing 1 mM glucose per minute at 28 °C. Cellulase was measured using 1% (w/v) carboxymethyl cellulose (CMC) in phosphate buffer (pH 7.0); reducing sugars were quantified at 540 nm using glucose as the standard and reported as U mL⁻¹, where one unit corresponded to 1 mM glucose released per minute at 28 °C (Miller, 1959). Similarly, Pectinase activity was determined using 1% (w/v) citrus pectin in phosphate buffer (pH 7.0) as the substrate; one unit was the amount of enzyme releasing 1 mM glucose per minute at 28 °C (Guan et al., 2020). Xylanase activity according to (Shi et al., 2011) using 1% (w/v) birchwood xylan in phosphate buffer (pH 7.0); one unit was defined as the amount of enzyme releasing 1 mM glucose per minute at 28 °C. For all assays, reaction mixtures prepared with uninoculated production medium served as controls.

## Results

### Sequencing statistics

The culture-independent high-throughput Illumina paired-end sequencing of the life-stages (larval, pupal, and adult), different populations (lab-bred, wild collection) of the two *Ips* pine beetles, *I. sexdentatus* (ISX) and *I. acuminatus* (IAC), along with wood tissue (control wood, fed/gallery wood) samples generated 8,670,808 raw reads. A Phred Quality score >30 was used for quality checks to obtain 7,147,099 clean reads (Supplementary Excel 1). The rarefaction curve and Good’s coverage indicator (>99%) indicated the sequencing depth for all samples (Supplementary Table 4, Supplementary Figure 1). The sample description, including abbreviations used, is provided in Table 1.

**Table 1.** Sample description.

| Sample Code | Sample details | Collection |
| --- | --- | --- |
| Ctrl.W | Uninfested pine wood control for lab rearing | Under lab environment |
| Fed.W | *Ips sexdentatus* infested gallery wood from F2 generation | Under lab environment |
| ISX.Larvae | *Ips sexdentatus* larvae from F2 generation | Lab-bred |
| ISX.Pupae | *Ips sexdentatus* pupae from F2 generation | Lab- bred |
| ISX.Adult | *Ips sexdentatus* adults from F2 generation | Lab-bred |
| ISX.WL.Adult | *Ips sexdentatus* wild-collected adults | Collected from forest |
| ISX.Ctrl.W | Uninfested wood control for *Ips sexdentatus* | Collected from forest |
| ISX.Fed.W | *Ips sexdentatus* infested gallery wood | Collected from forest |
| IAC.Larvae | *Ips acuminatus* larvae from F2 generation | Lab-bred |
| IAC.Pupae | *Ips acuminatus* pupae from F2 generation | Lab-bred |
| IAC.Adult | *Ips acuminatus* adult from F2 generation | Lab-bred |
| IAC.WL.Adult | *Ips acuminatus* wild-collected adults | Collected from forest |
| IAC.Ctrl.W | Uninfested wood control for *Ips acuminatus* | Collected from forest |
| IAC.Fed.W | *Ips acuminatus* infested gallery or fed wood | Collected from forest |

### Mycobiome composition and diversity

#### Fungal taxonomic abundance

The fungal ITS2 sequences were categorised into 2506 ASVs representing 63 fungal orders (Supplementary Excel 2). The computed Goods coverage index (0.98), representing filtered sequence reads (n > 5), validates that most ASVs in the samples were detected, with about 2% remaining undetected during sequencing. The predominant fungal orders were Saccharomycetales (ISX.Larvae- 0.39±0.08, ISX.Pupae-0.55±0.17, ISX.Adult-0.79±0.06, IAC.Larvae-0.60±0.04, ISX.WL.Adult-0.44±0.06, IAC.WLA-0.64±0.07, IAC.Fed.W-0.51±0.16) and Ophiostomatales (IAC.Pupae-0.67±0.07, IAC.Adult-0.50±0.12, Fed.W-0.42±0.12, Ctrl.W-0.69±0.04, ISX.Fed.W- 0.44±0.01) for all beetle and wood samples except in ISX.Ctrl.W, where Helotiales (0.48±0.03) was the dominant order (Figure 1A, Supplementary Excel 3). The heatmap depicting the top 35 fungal genera in different life stages of both beetles showed varied dominance of fungal genera at specific life stages (Figure 1B). For instance, in the case of ISX beetle life stages, the larvae were dominated by *the Talaromyces, Clonostachys, Acremonium, and Paratritirachium* genera. While *Atractiella*, *Candida*, *and Cladosporium* were the predominant fungal genera in pupae, *Cryptococcus* and *Wickerhamomyces* were prevalent in adults. Similarly, in IAC life stages, a prevalence of *Kuraishia* and *Nakazawaea* was observed in IAC larvae, while *Graphilbum* and *Aspergillus* were the dominant species in adults.

**Figure 1.**
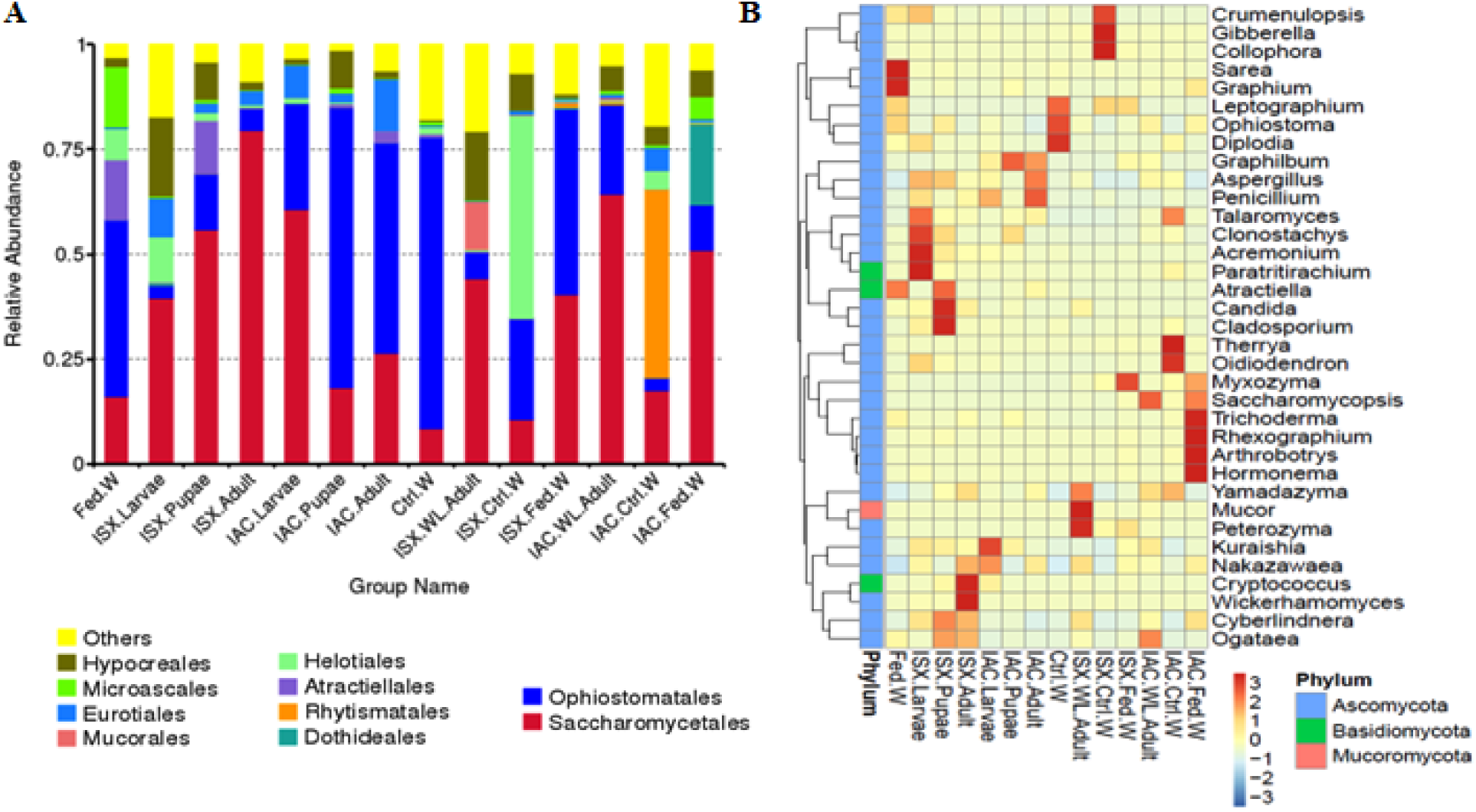
Fungal diversity in life stages (lab-bred), wild-collected adult pine-feeding beetles, and wood samples. (A) Bar diagram depicting beetle fungal community relative abundance at the order level (top 10). (B) 35 dominant fungal genera among life stages (lab-bred), wild adult, and wood samples are represented in the heatmap. The color gradient represents the relative abundance of ASVs, where light color represents lower abundance and darker color indicates higher abundance for a specific fungal genus. IAC- *I. acuminatus*; ISX- *I. sexdentatus*; WL- wild-collected; Ctrl. W- Control uninfested wood; Fed. W- Gallery wood.

#### ASV abundance

The ASV analysis revealed a diverse distribution of ASVs across different developmental stages for both *Ips* beetles. In the *I. acuminatus* (IAC), the core fungal communities across the life stages comprised of 47 ASVs, categorised into 16 families and 20 genera, with the predominance of Aspergillaceae, Ophiostomataceae, Trichocomaceae, Saccharomycetales_fam_Incertae_sedis families and fungal genera including *Penicillium*, *Graphilbum*, *Ophiostoma*, *Talaromyces*, *Kuraishia* (Figure 2A, Supplementary Excel 4). However, it is essential to note that each ASV may not represent a distinct species.

**Figure 2.**
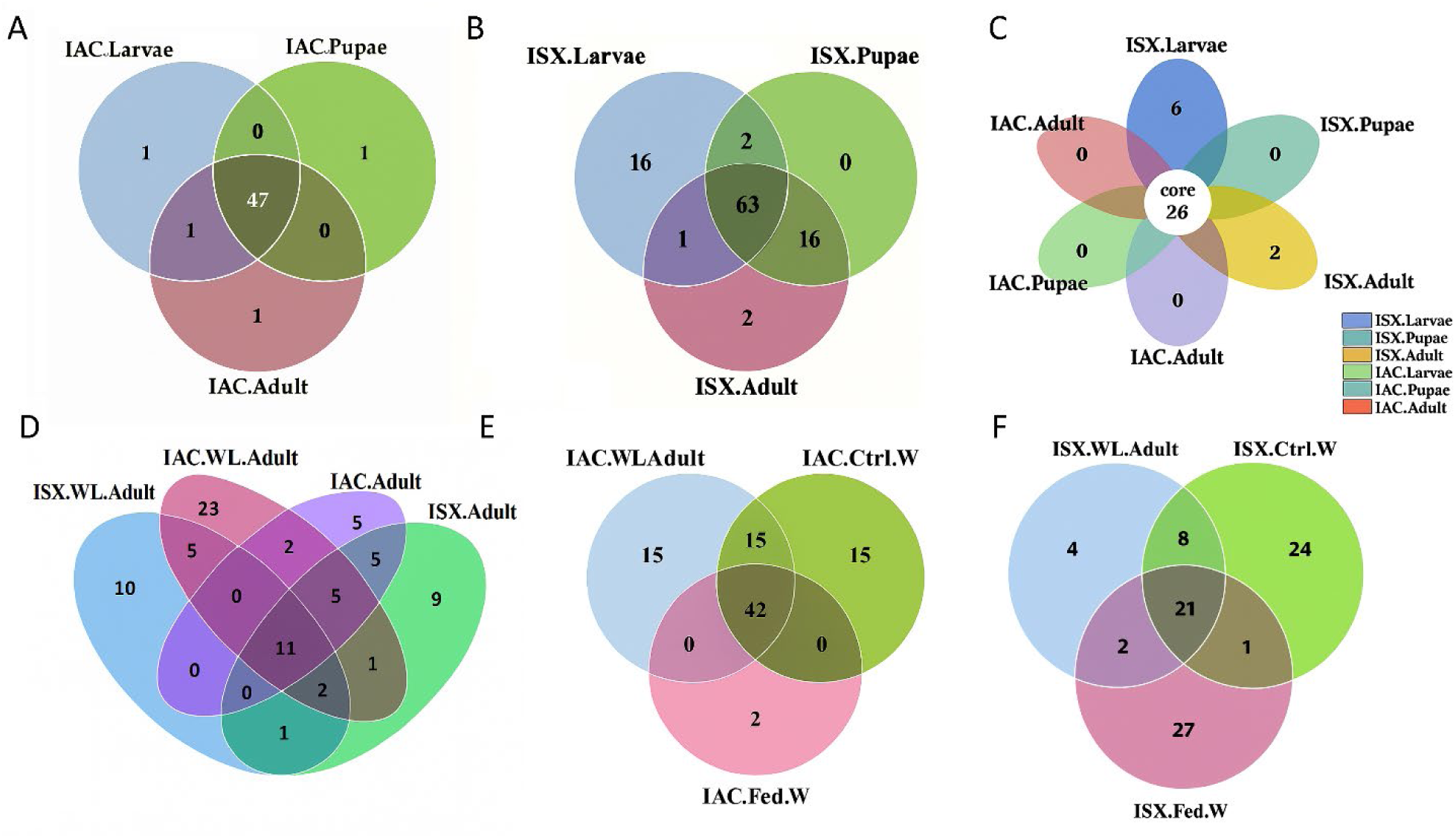
Mycobiome composition in life stages (lab-bred), wild adult pine-feeding beetles, and wood samples. (A) Venn diagram representing fungal ASV distribution in *I. acuminatus* life stages (IAC.Larvae, IAC.Pupae, IAC.Adult). (B) Venn diagram illustrating fungal ASV distribution in *I. sexdentatus* life stages (ISX.Larvae, ISX.Pupae, ISX.Adult). (C) The core ASVs across the life stages of two pine beetles are depicted by flower diagram. (D) The shared and unique ASVs present in *I. acuminatus, I. sexdentatus* wild beetles (IAC.WL.Adult, ISX.WL.Adult), and lab-bred beetles (IAC.Adult, ISX.Adult) are represented in the Venn diagram. (E) The fungal ASVs contribution of the wood mycobiome in shaping *I. acuminatus* mycobiome is shown in the Venn diagram. (F) The fungal ASVs contribution of wood mycobiome in shaping *I. sexdentatus* mycobiome is shown in the Venn diagram.

The core mycobiome shared between *I. sexdentatus* (ISX) life stages was represented by 63 ASVs, which were differentiated into 26 families and 32 genera (Figure 2B, Supplementary Excel 5). Highly abundant fungal families included Saccharomycetales_fam_Incertae_sedis, Pichiaceae, Ophiostomataceae, and fungal genera such as *Kuraishia*, *Ogataea*, *Ophiostoma*, and *Graphilbum* were prevalent. The ASV abundance data also revealed the core fungal mycobiome, including all life stages (larvae, pupae, adults) of both *Ips* pine beetles, which accounted for 26 ASVs categorised into 14 families and 16 genera (Figure 2C, Supplementary Excel 6). The core mycobiome comprised highly abundant fungal genera, including *Nakazawaea*, *Ogataea*, *Ophiostoma*, *Graphilbum*, *Kuraishia*, and *Talaromyces*. This pattern, where the number of shared ASVs is higher than individual stage-specific ASV abundance, suggests a conserved mycobiome structure across developmental stages within each species, with a stable set of core fungal taxa persisting throughout host development, despite stage-specific shifts in abundance.

When comparing the lab-bred and wild-collected beetles in both *Ips* species, the number of unique ASVs was higher than the shared ones, indicating that environmental factors influence the shaping of the mycobiome composition (Figure 2D, Supplementary Excel 7). In the case of IAC wild-collected beetles, the unique ASVs comprised 4 families and 5 genera (23 ASVs), while in IAC lab-bred adults, it was 2 families and 2 genera (5 unique ASVs). Similarly, for ISX, wild-collected and lab-bred beetles had a higher number of unique ASVs (10 and 9, respectively) compared to one shared ASV. The unique ASVs in wild-collected beetles belonged to 7 families and 7 genera, whereas in lab-bred beetles, they were found in 4 families and 4 genera. Beetles collected from the wild had a higher number of unique ASVs and greater diversity than those raised in the lab, reflecting mycobial enrichment while in natural habitats. Surprisingly, in both species, the common ASV between wild-collected and lab-bred belongs to the genus *Ophiostoma*, suggesting a conserved and potentially essential symbiotic association that persists irrespective of rearing conditions.

For both beetle species, wild-collected beetles shared more ASVs with control wood samples (IAC.WL.Adult and IAC.Ctrl.W-15 ASVs, ISX.WL.Adult and ISX.Ctrl.W-8ASVs) rather than with fed (gallery) wood samples (IAC.WL.Adult and IAC.Fed.W-0 ASVs, ISX.WL.Adult and ISX.Fed.W-2 ASVs). Subsequently, IAC control and fed wood had 15 and 2 unique ASVs, respectively. Similarly, ISX control and fed wood samples had respective 24 and 27 unique ASVs (Figures 2E and 2F, Supplementary Excel 8). These patterns suggest that beetle feeding induces a marked shift in the wood mycobiome.

#### Alpha diversity

The fungal mycobiome diversity was studied using alpha diversity indices (community richness, evenness, and diversity), revealing significantly different developmental stage-specific fungal associations. In *I. acuminatus* developmental stages, all the alpha diversity indices were higher in IAC.Larvae than IAC.Pupae and IAC.Adult samples (Figure 3, Supplementary Table 4, Supplementary Excel 9). IAC.Larvae (Pielou-0.45±0.06, Shannon-3.19±0.14, Simpson-0.79±0.02) and IAC.Adult samples (Pielou-0.35±0.02, Shannon-2.37±0.17, Simpson-0.61±0.05) exhibited significant differences (p<0.05) in fungal evenness and diversity. Similar to *I. acuminatus* life stages, in *I. sexdentatus* developmental stages, the fungal alpha diversity indices (community richness, evenness, diversity) were highest in ISX.Larvae than ISX.Pupae and ISX.Adult samples (Figure 3, Supplementary Table 4, Supplementary Excel 9). ISX.Larvae (Chao1: 308.84±45.22) samples showed significant differences (p<0.05) in community richness compared to the respective ISX.Pupae (Chao1-164.52±20.98) and ISX.Adult (Chao1-155.95±8.1) samples. Similarly, when comparing the larval samples of both species (IAC and ISX), significant differences (p < 0.05) in fungal richness were observed. The adult samples for both species also showed significant differences (p<0.01) in fungal richness and evenness. However, the pupal stage did not show any significant difference in community richness and evenness, ISX.Pupae (Shannon-3.7±0.33, Simpson-0.6±0.06) revealed significantly higher fungal diversity than IAC.Pupae (Shannon-2.55±0.29, Simpson-0.82±0.05) (p<0.05).

**Figure 3.**
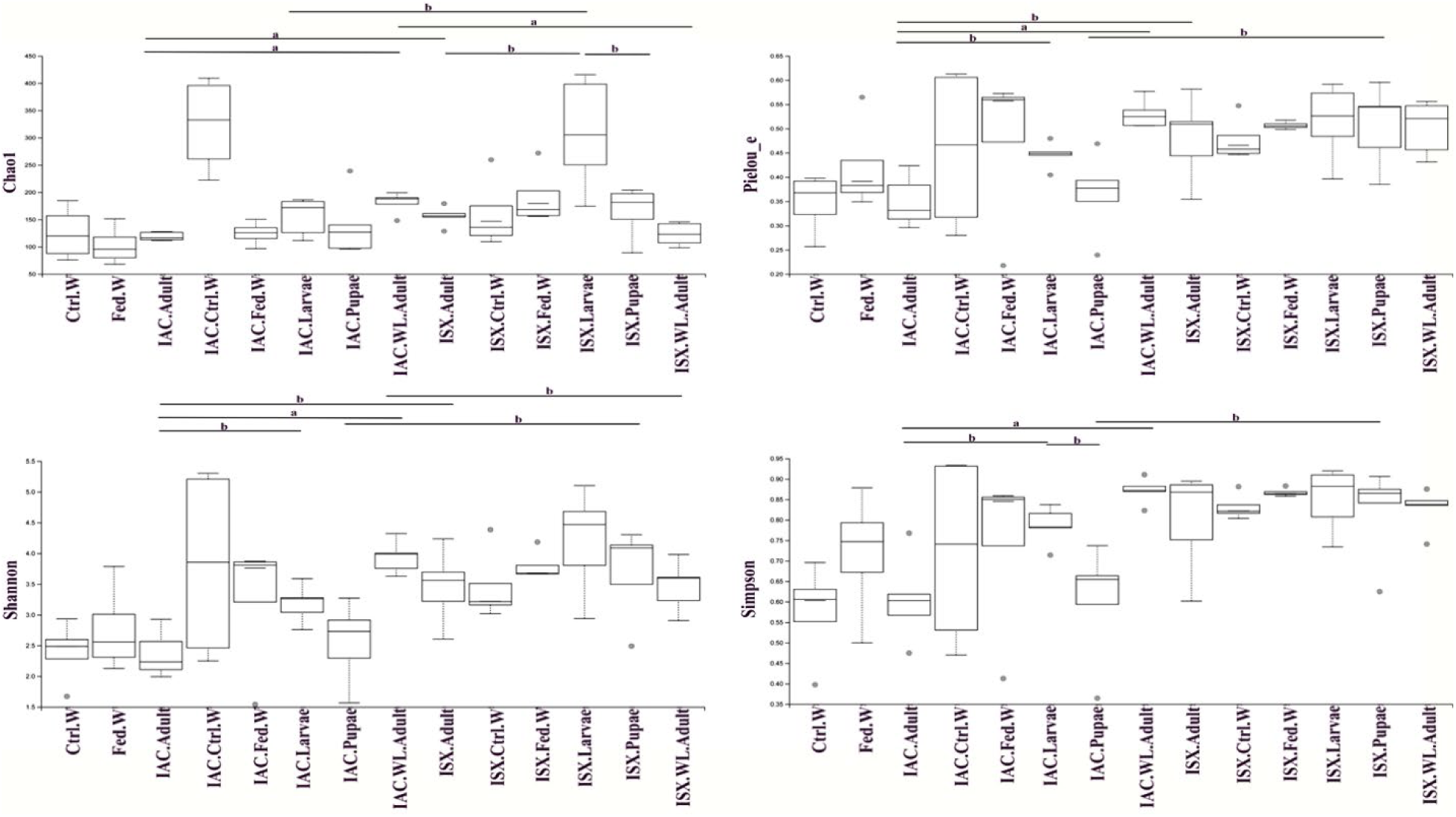
Alpha diversity indices depicted as boxplots for lab-bred, wild-collected *Ips* pine beetles and their wood tissue. (A) Fungal richness estimation (Chao1) depicts significant difference among different life stages, wild-collected adults, and wood samples. (B) Fungal evenness estimation by the Pielou index. (C) Shannon (D) Simpson index illustrates fungal diversity. Statistically significant differences among various groups at *p* <0.05 (designated as “b”), *p* <0.01 (designated as “a”) were observed using the Kruskal-wallis-pairwise-group test. IAC- *I. acuminatus*; ISX- *I. sexdentatus*; WL- wild-collected; Ctrl. W- Control uninfested wood; Fed. W- Gallery wood.

All the α-diversity indices showed significant differences (p<0.01) between wild-collected and lab-bred *I. acuminatus* beetles. However, such a significant difference was not observed between wild and lab-bred *I. sexdentatus* beetles. In contrast, studying the wood mycobiome at the IAC.Ctrl.W (324.13±45.29) samples showed significantly (p<0.05) higher fungal richness than their respective fed wood samples (124.69±11.07). Moreover, the fungal evenness and Shannon diversity index were higher in IAC.Ctrl.W samples, but the Simpson diversity index was higher in IAC.Fed.W samples. However, all the alpha diversity indices were higher in ISX.Fed.W samples compared to its control wood samples. Such observations suggest that beetle feeding affects the wood microbiome, and vice versa.

#### Beta diversity

The NMDS analysis, based on the unweighted UniFrac distance matrix, revealed clear separation of life stages (Figure 4). Notably, the fungal communities in the larvae of IAC and ISX exhibited significant differences both within and between groups. Furthermore, Metastat analysis indicated genus-specific shifts across the life stages of the pine beetle. Within *I. acuminatus*, larvae (IAC.Larvae) were enriched for *Kuraishia*, whereas adults (IAC.Adult) and pupae (IAC.Pupae) were dominated by *Graphilbum* (Table 2). A comparable pattern emerged in *I. sexdentatus* (ISX): *Talaromyces* predominated in larvae (ISX.Larvae), while *Ogataea* and *Cryptococcus* were more abundant in pupae and adults (ISX.Pupae, ISX.Adult). Wild-collected versus laboratory-bred contrasts further underscored these differences. In *I. acuminatus*, wild-collected beetles harboured significantly higher levels of *Kuraishia*, *Ogataea*, and *Ophiostoma*, whereas lab-bred beetles were enriched for *Graphilbum* and *Atractiella*. In *I. sexdentatus*, wild-collected beetles were characterised by significantly greater abundances of *Leptographium*, *Peterozyma*, and *Myxozyma*, while laboratory cohorts showed prevalence of *Kuraishia*, *Ogataea*, and *Wickerhamomyces*. Consistent presence was also evident in wood samples: IAC.Ctrl.W exhibited significantly higher abundances of *Therrya*, *Oidiodendron*, and *Talaromyces*, in contrast to IAC.Fed.W, where *Cyberlindnera*, *Ophiostoma*, and *Saccharomycopsis* predominated. In the ISX wood samples, ISX.Ctrl.W was dominated by *Crumenulopsis* and *Gibberella*, whereas *Graphilbum* and *Kuraishia* were abundant in ISX.Fed.W. Finally, multivariate testing (ADONIS and ANOSIM) confirmed significant compositional differences among life stages in both beetle species (Supplementary Tables 5, 6).

**Figure 4.**
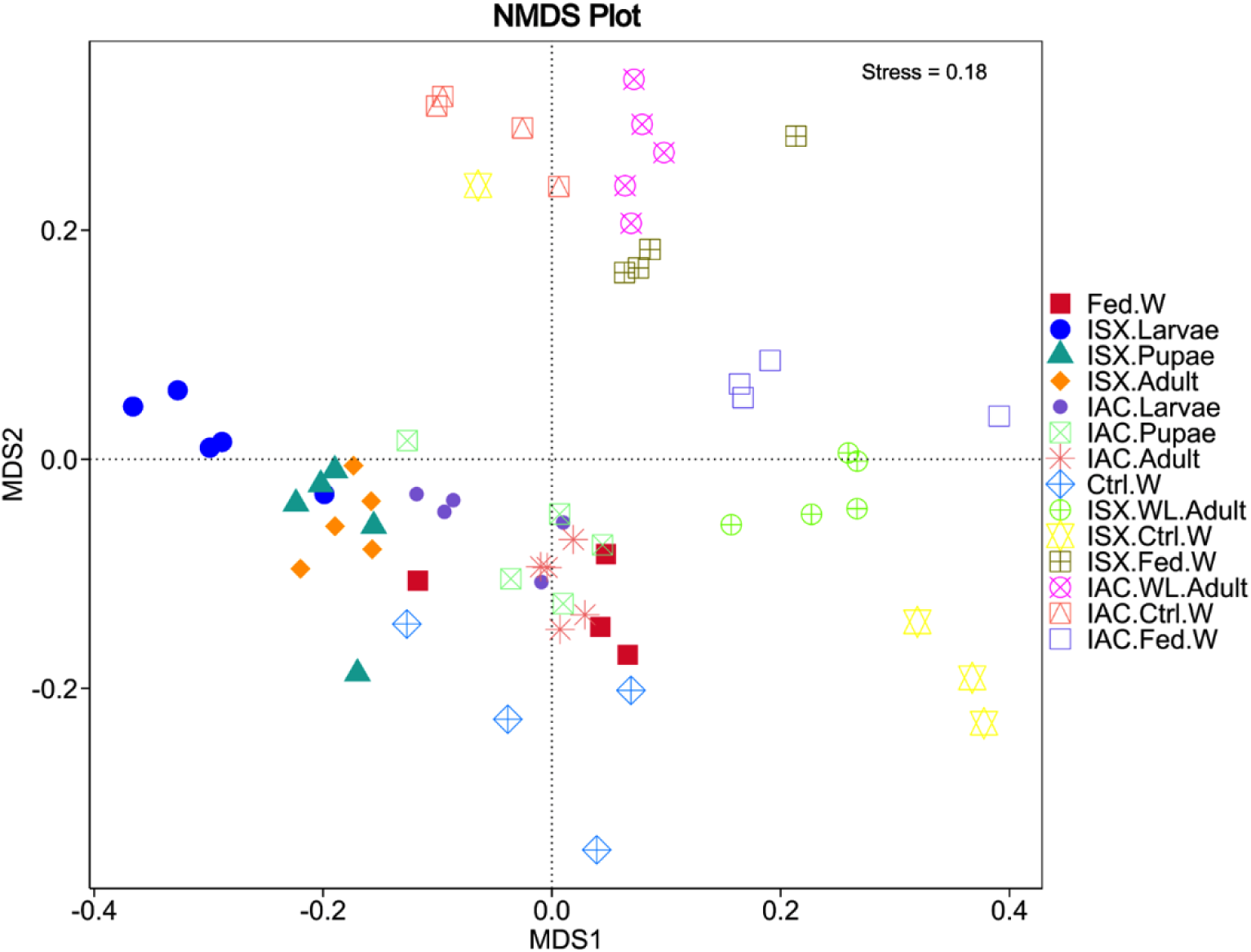
Beta diversity analysis. Non-metric Multi-Dimensional Scaling (NMDS) represents fungal diversity variation among life-stages and wild *Ips* pine beetles. IAC- *I. acuminatus*; ISX- *I. sexdentatus*; WL- wild-collected; Ctrl. W- Control uninfested wood; Fed. W- Gallery wood.

**Table 2.**
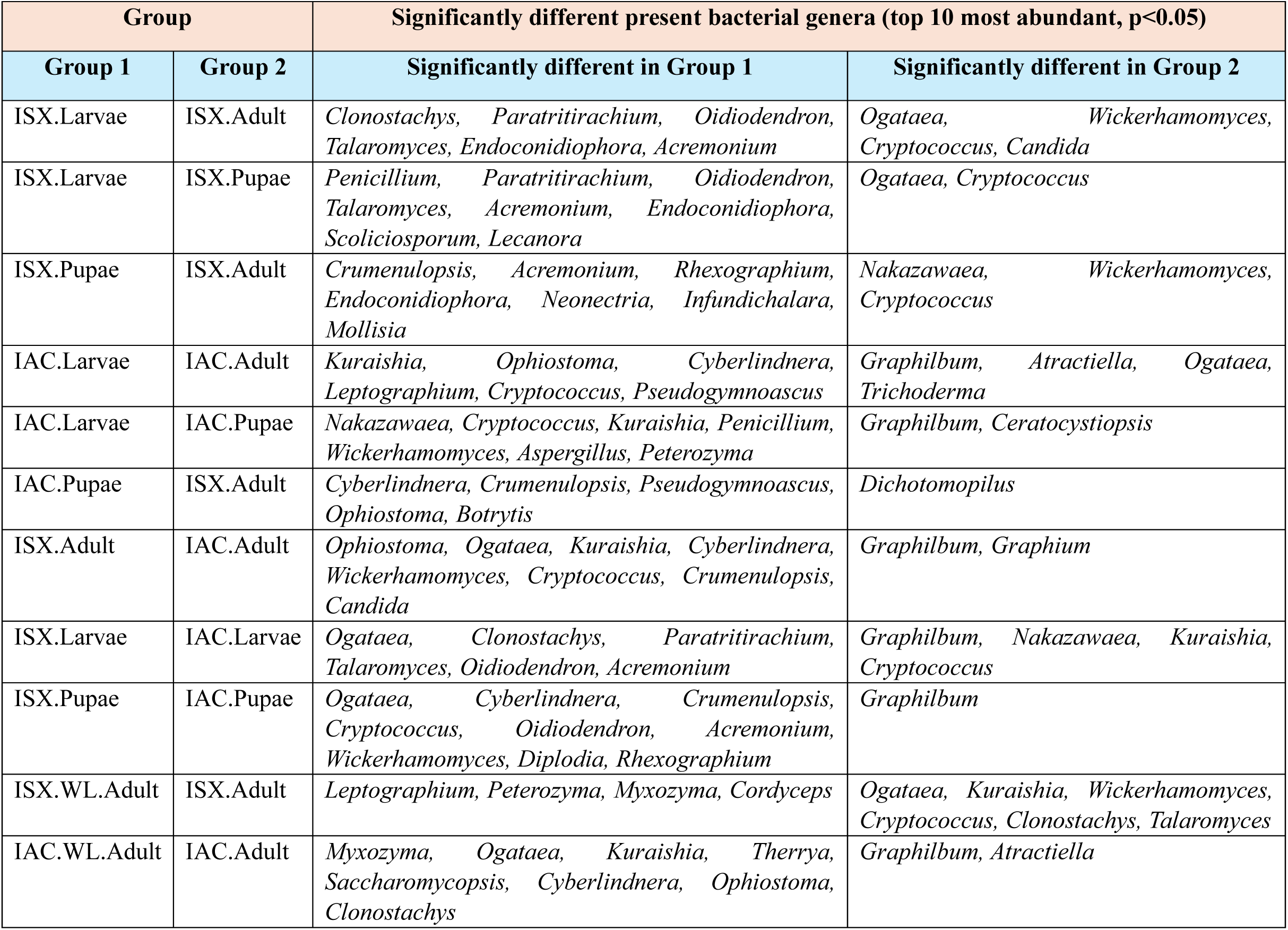
Metastat analysis revealing the top 10 differently abundant fungal genera across different developmental stages of *Ips* pine beetles.

Our study further investigates the presence of significantly dominant fungal genera across different life stages using LefSe analysis (Figure 5, Supplementary Figures 2, 3). In the case of *I. acuminatus*, distinct fungal families were identified as biomarkers at various life stages: the Hoehnelomycetaceae family (order: Tremellales) was prevalent in the IAC.Adult stage, while Ophiostomataceae (order: Ophiostomatales) was a key biomarker in IAC.Pupae. Additionally, Pichiaceae and Nectriaceae (order: Saccharomycetales) were dominant in IAC.Larvae (Figure 5A, Supplementary Figure 3A). Similarly, in *I. sexdentatus*, biomarkers varied across developmental stages: ISX.Larvae exhibited fungal biomarkers from the Helotiales and Tritirachiales orders, with Bionectriaceae and Tritirachiaceae families, while ISX.Pupae contained biomarkers from the Hoehnelomycetaceae family (order: Tremellales), and ISX.Adult was characterised by biomarkers from the Tremellaceae family (order: Tremellales) (Figure 5B, Supplementary Figure 3B).

**Figure 5.**
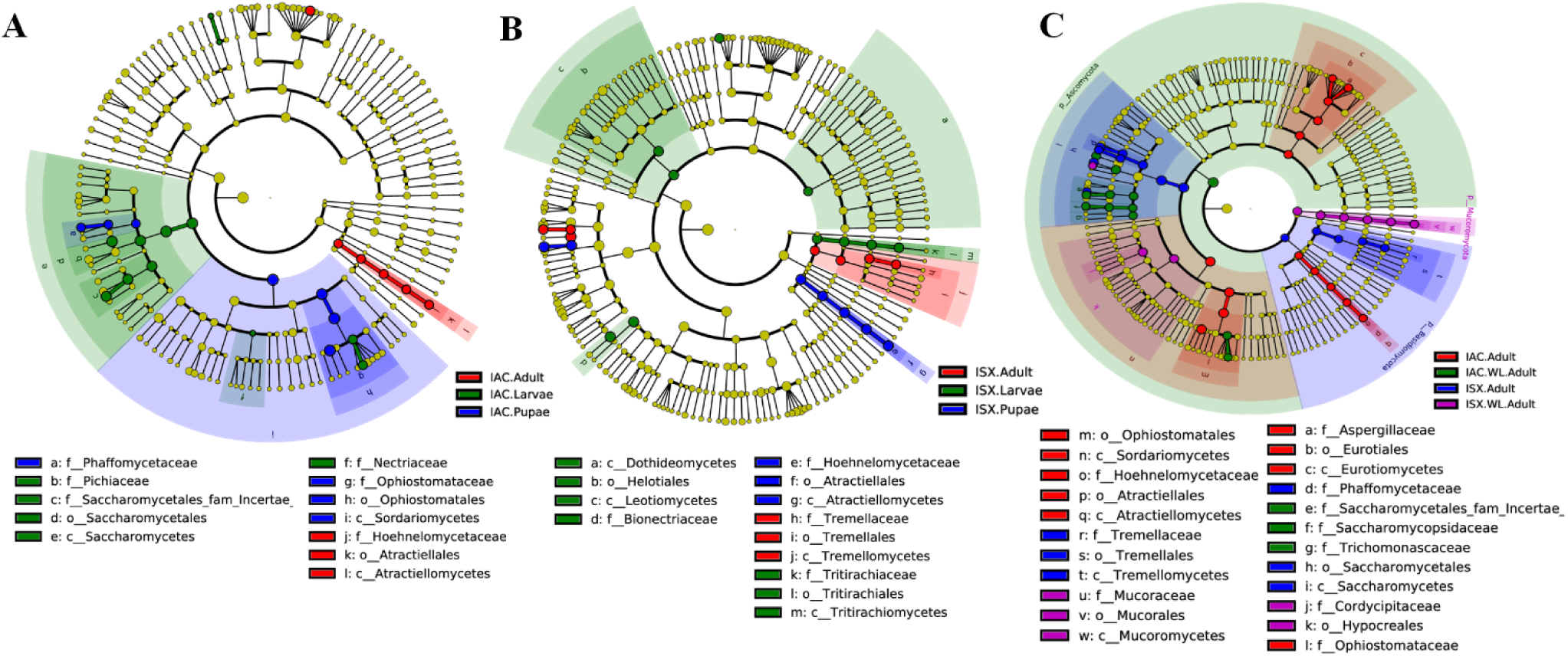
LefSe analysis. (A) Cladogram depicting significantly different fungal biomarkers across *I. acuminatus* life-stages. (B) Cladogram representing significantly distinct fungal biomarkers across *I. sexdentatus* life stages. (C) Cladogram depicting significant fungal biomarkers between wild-collected and lab-bred pine-feeding beetles. Taxonomic levels (phylum to genus) are outlined in the circle from inward to outward. The relative abundance of any taxon is portrayed by the size of the circles. Different coloured circles represent different life stages of the pine beetles. The letters above the different colored circles indicate a specific fungal biomarker. The yellowish-green circles illustrate non-significant fungal species. IAC- *I. acuminatus*; ISX- *I. sexdentatus*; WL- wild-collected.

When comparing lab-bred versus wild-collected beetles, distinct fungal biomarkers were observed in *I. acuminatus*: wild-collected beetles showed biomarkers from Saccharomycetales_fam_Incertae and Saccharomycopsideceae families, whereas lab-bred adults harboured biomarkers from Phaffomycetaceae and Tremellaceae families (orders: Saccharomycetales, Tremellales) (Figure 5C, Supplementary Figure 3C). Similarly, in *I. sexdentatus*, lab-bred beetles showed a dominance of Phaffomycetaceae and Tremellaceae families (orders: Saccharomycetales, Tremellales), while wild-collected specimens featured the Mucoraceae family (order: Mucorales) (Figure 5C, Supplementary Figure 3C).

Furthermore, distinct fungal biomarkers were identified across wild-collected beetles, control wood, and fed wood samples (Supplementary Figures 2, 3). In IAC.WL.Adult, the biomarkers included Cordycipitaceae and Ophiostomataceae families, while ISX.WL.Adult samples showed biomarkers from Phaffomycetaceae, Cordycipitaceae, and Mucoraceae families. For *I. acuminatus* wood samples, control wood contained Trichocomaceae, Myxotrichaceae, and Rhytismataceae families, whereas fed wood exhibited biomarkers from Dothioraceae and Hypocreaceae families. In *I. sexdentatus*, control wood (ISX.Ctrl.W) harboured Helotiales_fam_Incertae, while fed wood (ISX.Fed.W) showed Saccharomycetales_fam_Incertae as the dominant fungal biomarker.

### Functional prediction of beetle mycobiome

The putative roles of fungal communities across various life stages of both Ips species were assessed using FUNGuild (Figure 6). Both pine-feeding beetle species exhibited a prominent presence of saprotrophs, with notable variations in abundance, except for the pupal and adult stages of *I. acuminatus*. In the life stages of *I. sexdentatus*, saprotrophs were classified as wood saprotrophs, soil saprotrophs, and undefined saprotrophs. Notably, wild adults displayed a high prevalence of saprotrophs (72%), alongside a marked abundance of animal pathogens. Similarly, in *I. acuminatus*, the larvae (IAC.Larvae), pupae (IAC.Pupae), and wood-living adults (IAC.WL.Adult) exhibited a substantial presence of soil and wood saprotrophs.

**Figure 6.**
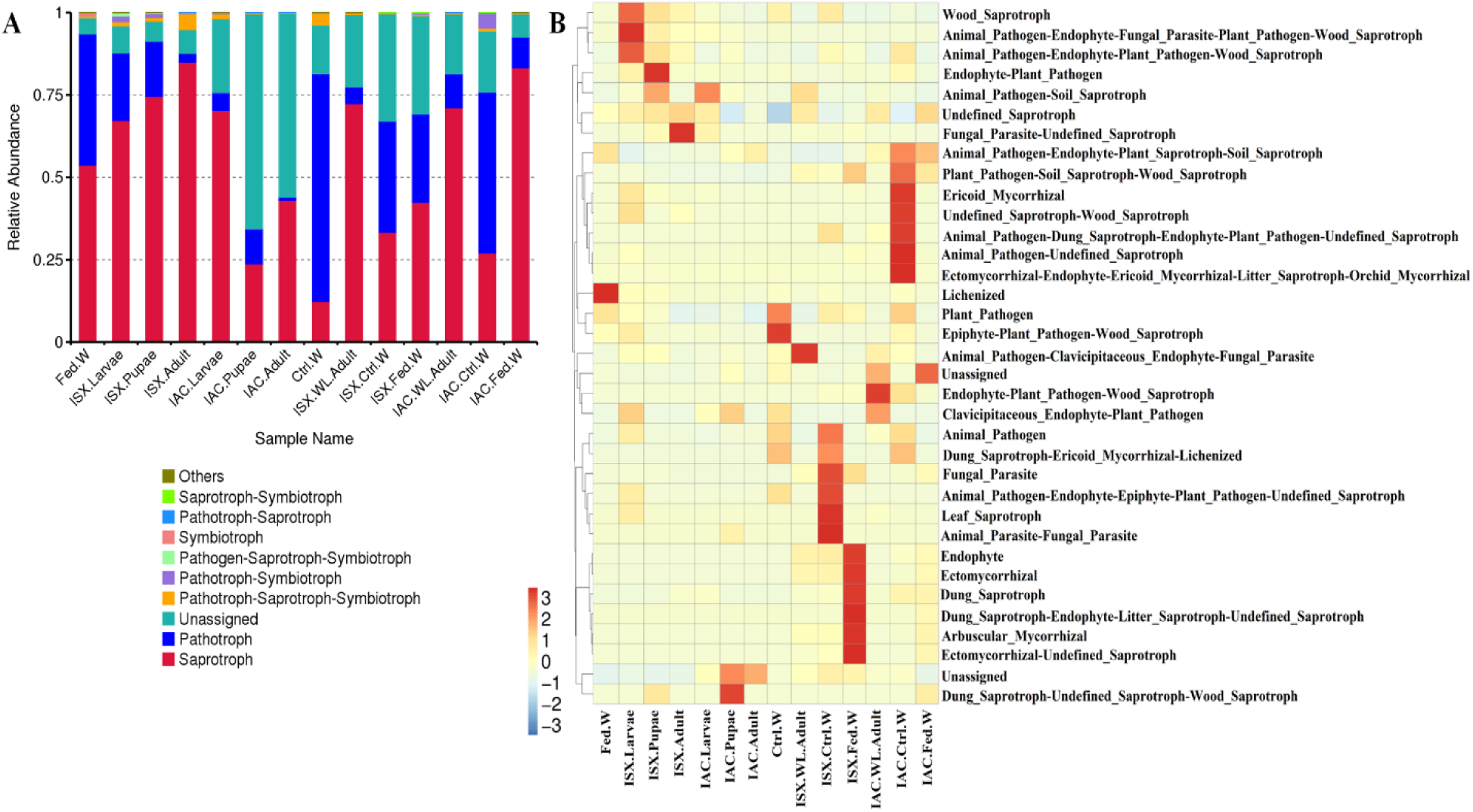
FUNGuild-based prediction of different functional groups (guilds) in the fungal community. (A) Bar diagram revealing the relative abundance of different fungal functional groups at the mode level. (B) Heatmap representing different fungal functional groups at the guild level. IAC- *I. acuminatus*; ISX- *I. sexdentatus*; WL- wild-collected; Ctrl. W- Control uninfested wood; Fed. W- Gallery wood.

### Fungal relative abundance determined by quantitative PCR assay

#### Ips sexdentatus life stages

qPCR assay revealed significant abundance differences in the total fungal population as well as selected fungal genera across the life stages in *I. sexdentatus*, which corroborated the metagenomic sequencing results (Figure 7, Supplementary Excel 10). The total fungal population in larvae was significantly higher compared to all other life stages (p < 0.01), while pupae and adults did not differ significantly. A similar pattern was observed for *Kuraishia*, where higher abundance was observed in larvae compared to adults and pupae (p < 0.01). For *Ophiostoma* and *Ogatea*, the pupae life-stage exhibited significantly higher abundance than adults (p < 0.05, p<0.01) and larvae (p < 0.05, p<0.01). However, for *Nakazawaea*, the adult exhibited significantly higher abundance than the larvae and pupae (p < 0.01 for both).

**Figure 7.**
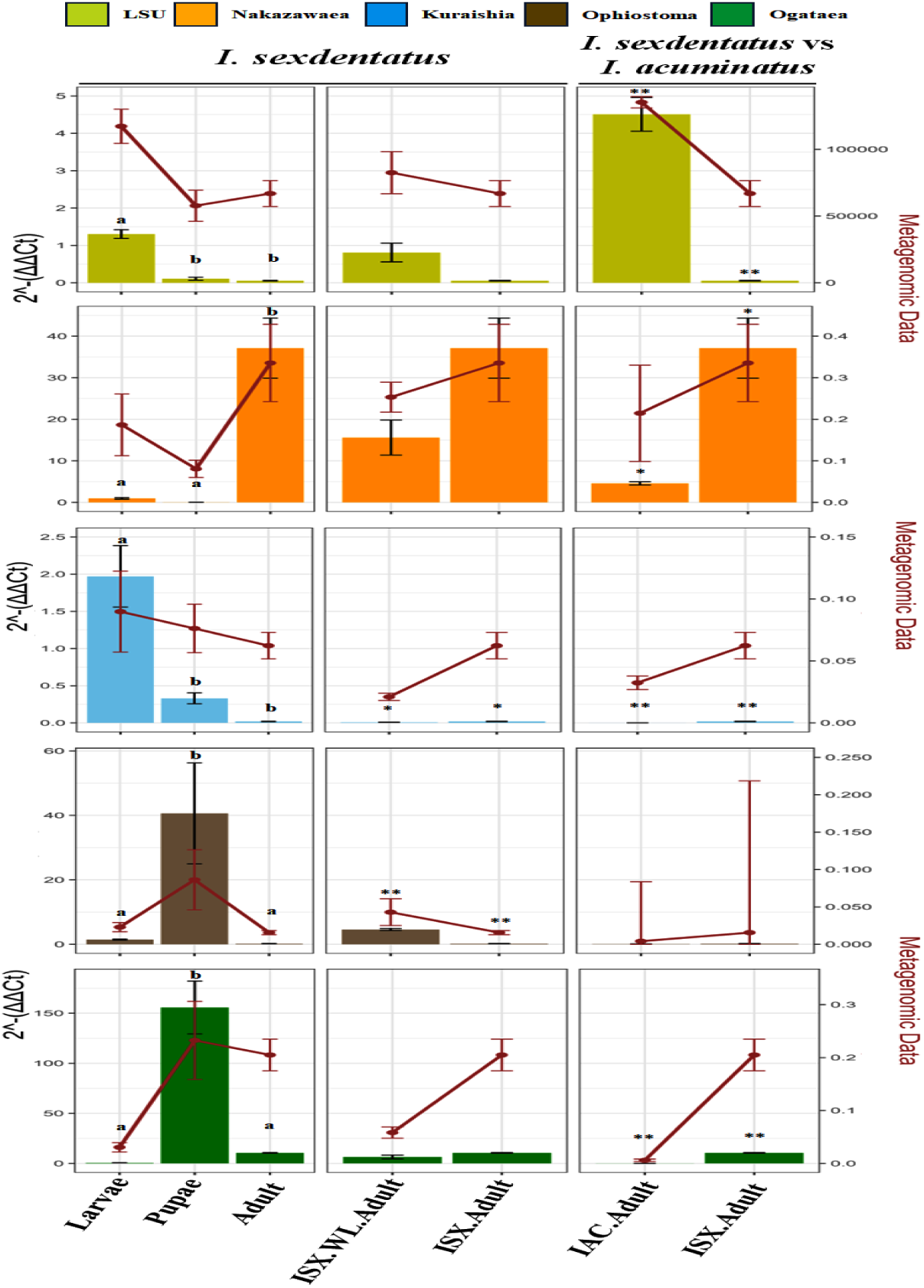
Quantitative PCR (qPCR) assays were performed to estimate the relative abundance of the predefined fungal taxa in the two pine-feeding beetle species. Fold change in fungal abundance was estimated using the 2−ΔΔCt method that was normalized against stable reference genes (ISX: β-Tubulin; IAC: EF1α and RPL7). Values are presented as means ± SE from beetle samples (n = 4). Statistical contrasts were determined using ANOVA, with post-hoc and t-test used for pair-wise comparisons. Different letters indicate differences between groups (p < 0.01). Full statistical results are presented in Supplementary Excel 10. IAC- *I. acuminatus*; ISX- *I. sexdentatus*; WL- wild-collected.

#### Wild-collected adults vs. lab-bred adults

Comparing wild-collected and lab-bred adults revealed a significant difference in the abundance of *Kuraishia* (p < 0.05) and *Ophiostoma* (p < 0.01) (Figure 7, Supplementary Excel 10). However, no such significant differences were observed for other genus-specific primers.

#### I. sexdenatatus vs. I. acuminatus adults

Comparisons between ISX *and* IAC adult beetles showed significant differences for all primers except *Ophiostoma* (Figure 7, Supplementary Excel 10). The relative abundance of the total fungal population was markedly higher in IAC adults (p < 0.01). However, for *Nakazawaea* (p < 0.05), *Kuraishia* (p < 0.05), and *Ogatea* (p < 0.01), their relative abundances were significantly higher in ISX adults. These findings suggest that certain fungal associates are more prevalent in ISX populations, while others, such as *Ophiostoma*, are maintained at comparable levels regardless of geographic origin.

### Effects of monoterpenes on pine bark beetle-associated yeasts

The four yeasts associated with the pine bark beetles, *Y. mexicana, N. holstii, K. molischiana, and C. mississippiensis* showed distinct growth responses with individual monoterpene treatment as well as monoterpene blend (Figure 8). In the control treatments, consisting of growth media (PDB) with no supplements and growth media with DMSO, all isolates exhibited consistent growth and a sharp rise in turbidity at approximately 40 hours. Exposure to individual monoterpenes, including (−)-α-pinene, 3-carene, and terpinolene, caused species-specific growth reductions. The highest degree of growth inhibition due to monoterpenes was most frequently observed with (−)-α-pinene, particularly in *Y. mexicana* and *N. holstii*. In contrast, 3-carene and terpinolene caused moderate inhibition in most isolates. Notably, the growth curves indicated that inhibition was most pronounced during the first 20–24 hours, followed by a later recovery or lower inhibition, which suggested a possible acclimation period. The monoterpene blend (a mix of α-pinene, 3-carene, and terpinolene) that mimics the more complex terpene state encountered within infested host tissue was consistently and significantly most effective at inhibiting growth in all the yeast species.

**Figure 8.**
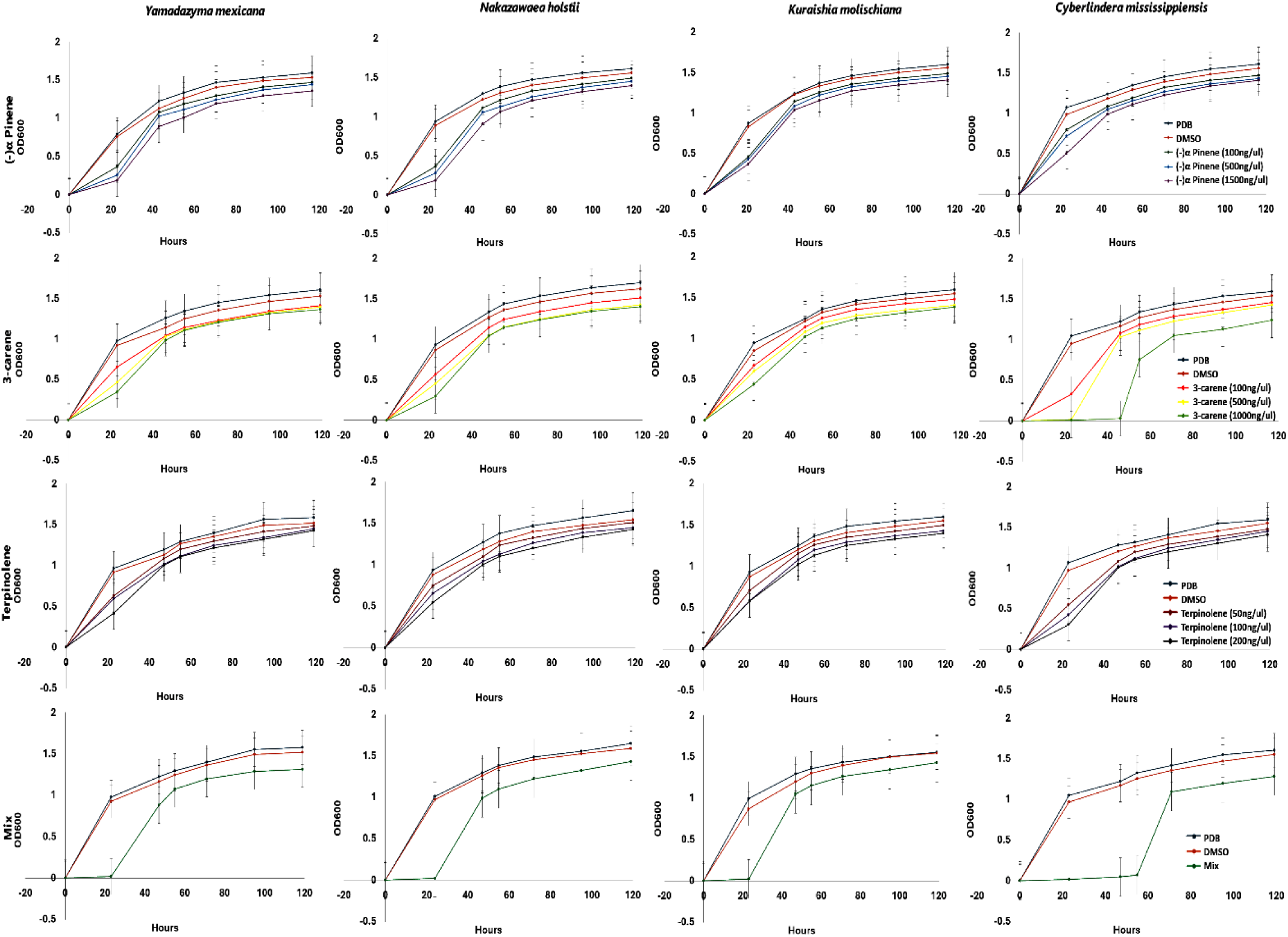
Bioassay of yeast isolates subjected to monoterpenes. Growth response of four yeast isolates (*Yamadazyma mexicana, Nakazawaea holstii, Kuraishia molischiana,* and *Cyberlindera mississippiensis*) to different monoterpenes: α-pinene, 3-carene, terpinolene, and their mixture.

### Gut microbial colonisation and biofilm formation

The symbiotic association in the different gut regions of *I. sexdentatus* was observed through an SEM study (Supplementary Figure 4). Colonisation of the yeast in the specific gut regions, i.e., midgut (Supplementary Figure 4A, 4B) and hindgut (Supplementary Figure 4C, 4D), is very prominent and significant. Similarly, *in vitro* biofilm formation by the gut isolates was observed under SEM, indicating the adhesion of yeast cells to the experimental surfaces. In addition, these gut isolates exhibited multi-layered, three-dimensional co-aggregation and a tendency to attach to solid surfaces (Supplementary Figure 5).

### Fungal interactions

Antifungal potential of the four yeast isolates was observed under SEM, where prominent degradation and deformation of the selected EPF and NEPF mycelial structures were visualised. The healthy mycelial structure of the EPF (Figure 9A-D) as well as NEPF (Figure 10A, F), and individual yeast cellular morphology (Supplementary Figure 6) were also documented through SEM. The antagonistic effect of all four gut symbionts on the EPF, e.g., *A. caatinguens* (Figure 9 E-H), *B. bassiana* (Figure 9I-L), *C. rosea* (Figure 9M-P), and *Trichoderma* sp. (Figure 9Q-T), was reflected by a massive deconstruction of the EPF mycelial structure along with the interactive yeast cells, specifically on the dismantled mycelial morphologies in comparison to the healthy fungal mycelia. Interestingly, the yeast symbionts were unable to make prominent disorientation on the NEPF mycelial structures of *Ophiostomatoid hongxingense* (Figure 10B-E) and *Ophiostomatoid piceae* (Figure 10G-J) in comparison to the EPF.

**Figure 9.**
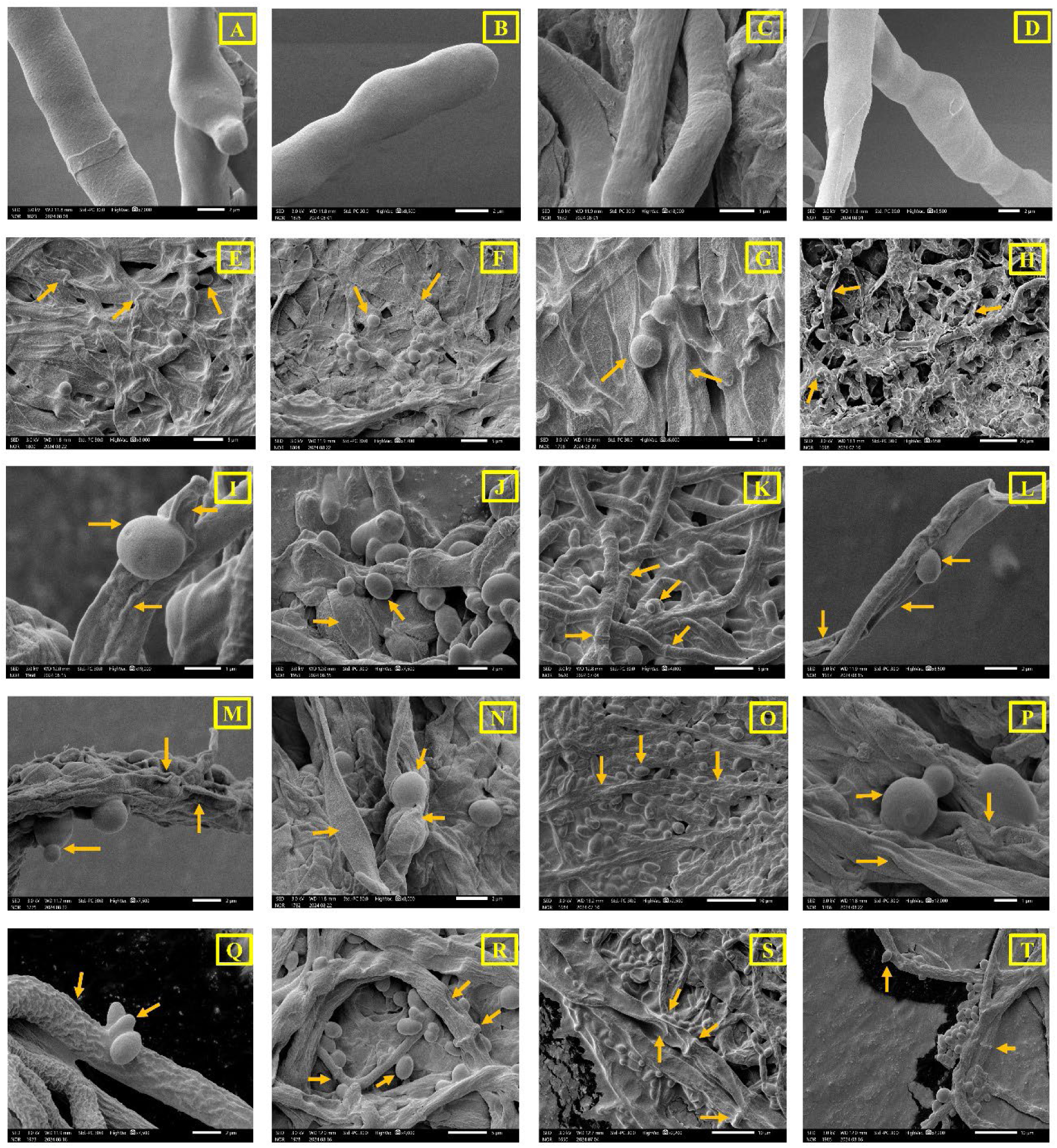
Interaction of symbiotic yeasts against entomopathogenic pathogenic fungi. (A-D) Healthy fungal mycelial structure of entomopathogenic fungi [(A) *Absidia caatinguens* (B) *Beauvaria bassiana* (C) *Clonostachys rosea* (D) *Trichoderma sp*. (E-H) SEM observation between entomopathogenic fungi *Absidia caatinguens* and yeasts isolates [(E) *N. holstii* (F) *C. missipplensis* (G) *Y. mexicana* (H) *K. molischiana*]. (I-L) SEM observation between entomopathogenic fungi *Beauveria bassiana* and yeast isolates [(I) *N. holstii* (J) *C. missipplensis* (K) *Y. mexicana* (L) *K. molischiana*]. (M-P) SEM observation between entomopathogenic fungi *Clonostachys rosea* and yeast isolates [(M) *N. holstii* (N) *C. missipplensis* (O) *Y. mexicana* (P) *K. molischiana*]. (Q-T) SEM observation between entomopathogenic fungi *Absidia caatinguens* and Symbiotic yeast [(Q) *N. holstii* (R) *C. missipplensis* (S) *Y. mexicana* (T) *K. molischiana*]. Yellow arrows indicated different magnitudes of entomopathogenic fungal hyphae damage.

**Figure 10.**
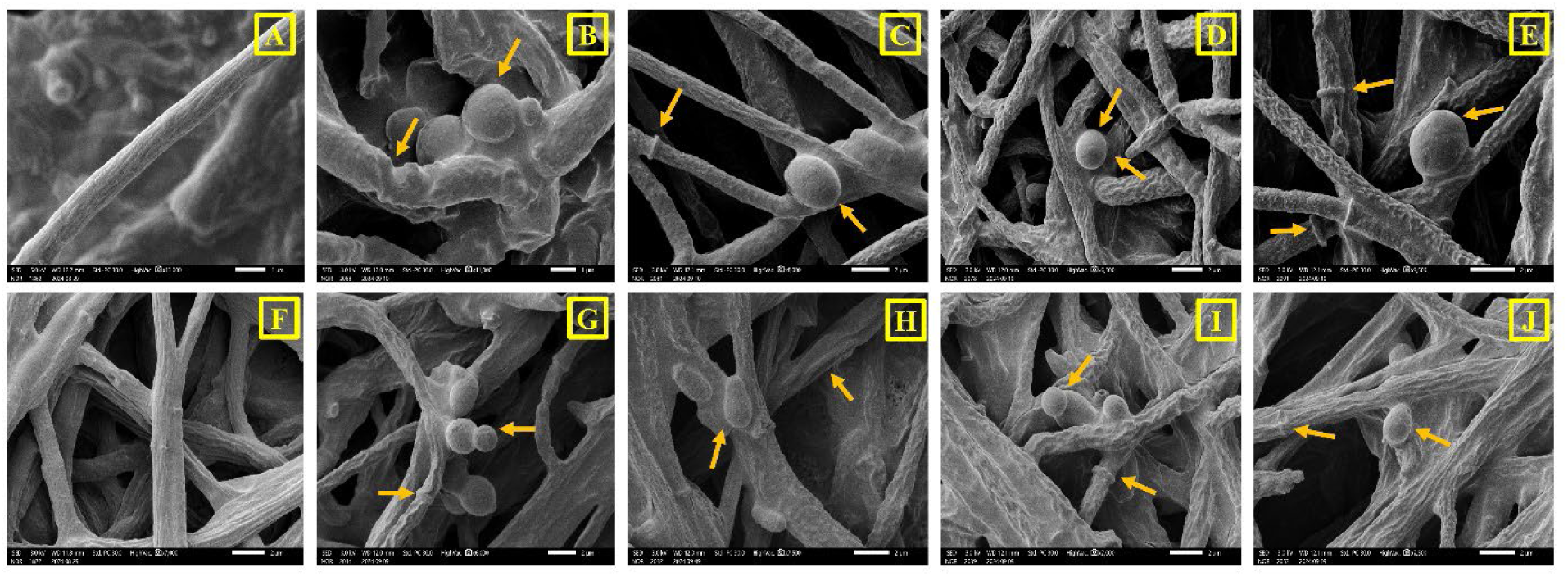
Interaction of symbiotic yeasts against non-entomopathogenic fungi. (A) Healthy fungal mycelial structure of non-entomopathogenic fungi *Ophiostoma hongxingense*. (B-E) SEM observation between non-entomopathogenic fungi *Ophiostoma hongxingense* and symbiotic yeast [(B) *N. holstii* (C) *C. missipplensis* (D) *Y. mexicana* (E) *K. molischiana*]. (F) Healthy mycelial structure of *Ophiostoma piceae.* (G-J) SEM observation between non-entomopathogenic fungi *Ophiostoma multisynnematum* and yeast isolates [(G) *N. holstii* (H) *C. missipplensis* (I) *Y. mexicana* (J) *K. molischiana*].

### Enzyme production

All four yeast strains were assessed for their antifungal activity and digestive enzymatic potential using a qualitative index (QI). The isolated yeast strains exhibited varying degrees of hydrolytic potential against starch, laminarin, cellulose, pectin, and xylan, as indicated by the corresponding qualitative index shown in Figure 11A. Notable halo zone formation was observed for starch, cellulose, and pectin hydrolysis, as well as for other tested substrates (Figure 11B-E). No QI was detected for protein or chitin metabolism. Among the yeasts, *N. holstii* demonstrated activity only against starch and xylan hydrolysis. In contrast, *C. mississippiensis* exhibited active QI against all substrates. Additionally, the gut isolate *Y. mexicana* displayed QI for laminarin, cellulose, pectin, and xylan, while *K. molischiana* showed QI activity against cellulose, pectin, and xylan.

**Figure 11.**
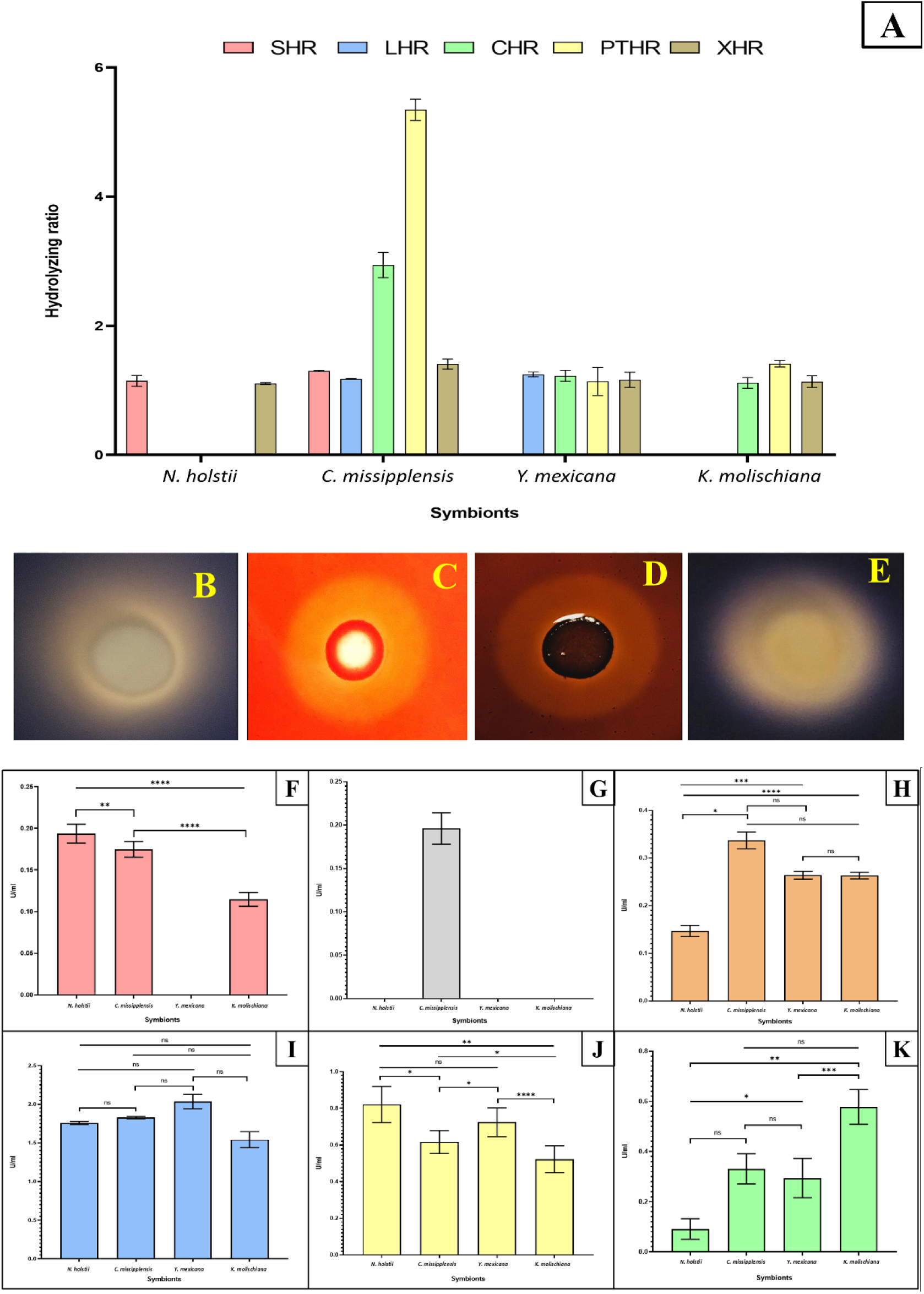
Detection of physiochemical potential of symbiotic yeasts. (A) Detection of the potential of symbiotic yeast in different antifungal and digestive enzyme production. SHR- Starch hydrolyzing potential, LHR- Laminarin hydrolyzing potential, CHR- Cellulose hydrolyzing potential, PTHR- Pectin hydrolyzing potential, XHR- Xylan hydrolyzing potential. Hydrolyzing zone forming potential of yeast symbionts. (B) Starch hydrolyzing capability of *Nakazawaea holstii*. (C) Starch hydrolyzing capability of *Cyberlindera missipplensis*. (D) Cellulose hydrolyzing ability of *C. missipplensis*. (E) Pectin hydrolyzing efficiency of *C. missipplensis*. Antifungal and digestive enzyme production capability of symbiotic yeast (F-K). (F) Amylase production capability of the yeast symbionts. (G) β-glucanase production capability of the yeast symbionts. (H) Cellulase production capability of the yeast symbionts. (I) Chitinase production capability of yeast symbionts. (J) Pectinase production capability of the yeast symbionts. (K) Xylanase production capability of the yeast symbionts. The standard error of the mean was calculated from three determinations. The significance level of the variables was determined by one-way ANOVA followed by Tukey test at a 95% confidence level (P < 0.05) using GraphPad Prism software (version 10.4.2). ** P value <0.05, *** P value <0.001, **** P value <0.0001, ns- non-significant.

### Antifungal and Digestive enzyme production

The four yeast isolates showed varying potential for antifungal and digestive enzyme production (Figure 11F-K). The amylase production capabilities of the gut symbionts *N. holstii*, *C. mississippiensis*, and *K. molischiana* were quantified as 0.193 ± 0.11, 0.175 ± 0.01, and 0.114 ± 0.08 U/ml, respectively (Figure 11F). Notably, β-glucanase production was observed exclusively in *C. mississippiensis* (0.19 ± 0.02 U/ml) (Figure 11G). Similarly, the cellulolytic efficiency of the yeast symbionts was assessed, revealing *C. mississippiensis* as the most prolific cellulase producer (0.33 ± 0.04 U/ml), followed by *K. molischiana* (0.26 ± 0.013 U/ml), *Y. mexicana* (0.25 ± 0.014 U/ml), and *N. holstii* (0.14 ± 0.01 U/ml) (Figure 11H). The yeast isolates *N. holstii*, *C. mississippiensis*, *Y. mexicana*, and *K. molischiana* demonstrated fungal cell wall lytic enzyme (Chitinase) production of 1.77 ± 0.03, 1.82 ± 0.03, 2.04 ± 0.08, and 1.56 ± 0.13 U/ml, respectively (Figure 11I). Furthermore, substantial pectinase production was observed by the four yeast isolates (*N. holstii-* 0.84±0.08; *C. mississippiensis*- 0.61±0.06; *Y. mexicana*- 0.72±0.08, and *K. molischiana* 0.53±0.07 U/ml) (Figure 11J). Additionally, the xylanase production potential of these symbionts was evaluated, with *K. molischiana* emerging as the most efficient producer (0.57 ± 0.07 U/ml), followed by *C. mississippiensis* (0.32 ± 0.06 U/ml), *Y. mexicana* (0.29 ± 0.08 U/ml), and *N. holstii* (0.08 ± 0.04 U/ml) (Figure 11K). However, none of the symbionts produced a significant quantity of protease.

## Discussion

The relationship between bark beetles and their associated fungi is intricate and dynamic, encompassing a range of interactions that can vary from mutualism and commensalism to antagonism, depending on the context (Hofstetter, 2015). These associations may shift from beneficial to harmful, influenced by the life stages of the beetles and changing environmental conditions (Six, 2021). To fully understand the context-dependent dynamics in *Ips* bark beetles, it is essential to explore the fungal communities associated with different developmental stages. Moreover, environmental factors and plant-insect interactions play a pivotal role in shaping the microbial communities of insects, including bark beetles (Baños-Quintana, 2024; Chakraborty, 2023). This study thus aims to investigate the impact of developmental stages and environmental conditions on the fungal community associated with two species of pine-feeding *Ips* bark beetles. We investigated the mycobiome of different developmental stages in pine beetles and their interactions. However, we did not resolve sex-specific microbiota in adult beetles. Sex-dependent variation in microbial associations and its functional significance, therefore, remains an open question.

Our findings reveal persistence of a core fungal community, including fungal genera such as *Kuraishia*, *Ogataea*, *Ophiostoma*, *Graphilbum*, and *Cyberlindnera*, across the developmental stages of two pine-feeding bark beetles, *Ips acuminatus* and *Ips sexdentatus*. The association of such fungal genera with other *Ips* bark beetles has been previously reported (Baños-Quintana, 2024; Chakraborty, 2020, 2023). The occurrence of a persistent fungal community across developmental stages indicates its pivotal role in supporting the host (Barcoto, 2020; Briones-Roblero, 2017). For instance, fungal genera such as *Kuraishia* and *Ogataea* can convert trans and cis-verbenol into anti-aggregation pheromone verbenone, providing a ‘microbe-mediated off-switch’ for beetle aggregation, regulating colony density, and reducing intraspecific competition. (Hunt, 1990). Additionally, the genome of *K. molischiana* reveals pathways for vitamin B6 and essential amino acids, which may benefit beetle nutrition (Cheng, 2023), while *Cyberlindnera* aids in detoxifying pine metabolites and degrading lipids and starch (Briones-Roblero, 2017). Similarly, other fungal associates, such as *Ophiostoma*, also have the ability to utilise plant defence compounds as carbon sources (Zaman, 2023). These findings highlight a metabolically adaptable mycobiome that supports host nutrition, detoxification, and chemical signalling, contributing to the beetles’ ecological success. Moreover, our study found greater fungal richness and diversity in gregarious larval stages, suggesting fungal acquisition aids nutrient assimilation and detoxification (Liu et al., 2022). Fungal diversity declined in the pupal stage, likely due to metamorphic changes (Sela et al., 2020), a pattern also observed in bacterial diversity (Khara et al., 2024; Peral-Aranega et al., 2023). Notably, *I. sexdentatus* exhibited higher fungal richness and diversity than *I. acuminatus*, reflecting differences in their feeding behaviors. While both species feed on the same pine, *I. acuminatus* targets thin-barked phloem, whereas *I. sexdentatus* feeds on thicker bark. Additionally, *I. acuminatus* clogs maternal tunnels with frass, promoting mutualistic fungus growth in thinner bark areas (Biedermann, 2020), resulting in distinct microhabitats that likely influence the fungal communities associated with each beetle.

Laboratory adaptation and breeding conditions significantly influence fungal associations, often with unpredictable outcomes (Augustinos, 2019; Ibarra-Juarez, 2018). In the wild, beetles face diverse environmental pressures that foster richer fungal communities, while controlled lab conditions may induce a bottleneck effect, limiting diversity (Augustinos, 2019). Wild-caught *I. acuminatus* exhibit greater fungal richness and diversity compared to lab-bred counterparts. However, similar data for *I. sexdentatus* are lacking, warranting further investigation into the ecological drivers behind these differences. Additionally, variations in collection years in our study may reflect the influence of differing environmental conditions.

It is well-documented that insects can acquire microbial communities horizontally as they consume their diet (Hartmann, 2017; Kikuchi, 2007; Sugio, 2015). Thus, it is expected that the host microbiome (in this case, wood) will influence the beetle microbiome. Our findings reveal that beetles share more amplicon sequence variants (ASVs) with control wood than with gallery wood. This higher sharing with control wood (unfed phloem) suggests microbial acquisition from the wood microbiome during feeding. This phenomenon is particularly notable for *Ips acuminatus* beetles, which feed on wood-associated fungi for nutritional benefits (Papek, 2024). Consequently, differences in the fungal community between fed and unfed wood may reflect changes in the fungal composition driven by beetle feeding. We hypothesise that during wood consumption, bark beetles introduce microbes into the gallery wood microbiome via oral secretions, exoskeleton contact, and faecal deposition, thereby altering the mycobiome. However, further research is required to fully understand the ecological implications of pine microbiome acquisition and exchange in these beetles.

FUNGuild-based functional predictions of the beetle-associated fungal community identified several putative functional groups. The consistent prevalence of saprotrophs across developmental stages highlights their role in facilitating beetle nutrition within a saprophytic environment. These fungi specialise in decomposing decaying organic matter, converting it into usable forms for beetles. Dominant genera, such as *Graphilbum*, *Leptographium*, and *Cryptococcus*, are well-known saprotrophs that derive nourishment from decomposed organic material, including fallen wood (Gladieux, 2011; Trollip, 2021). Nevertheless, it’s crucial to acknowledge that FUNGuild is a software that predicts functional abundances based on marker genes. Therefore, the putative functional prediction of the microbiome requires additional validation through metatranscriptomic, metaproteomic, or other functional assays.

The contrasting growth responses documented here underscore the importance of evaluating bark beetle-associated microbe growth in response to exposure to multiple terpene compounds. While studies tend to challenge microbial resistance against a single terpene, in the field, these microorganisms inside beetles face a dynamic blend of host monoterpenes that are emitted collectively upon resin flow following beetle attack (Adams, 2011; Chiu, 2017). Our results show that the monoterpene blend of α-pinene, 3-carene, and terpinolene consistently inhibited growth more than any of the individual compounds, suggesting the possibility of additive or even synergistic action of the combined terpenes. These interactions may arise from overlapping toxicity mechanisms, membrane disruption, or metabolic burden due to the need to detoxify multiple compounds simultaneously (Marmulla, 2014; Sittichok, 2025). The early inhibition of *Y. mexicana* and *N. holstii*, followed by partial recovery, indicates that these species may adapt over time but face a competitive disadvantage in high-terpene environments shortly after host colonisation. Future studies combining bioassays with metabolomics and transcriptomic analysis of detoxification pathways could provide deeper insights into terpene tolerance and the ecological roles of these yeasts in bark beetle microbiomes. Overall, exposure to multiple terpenes is crucial and should be conducted more frequently, as it more accurately reflects the selective pressures in Nature that shape bark beetle-associated microbial communities.

Microscopic imaging of gut colonisation, together with biofilm assays on culturable isolates, indicates that microbial assemblages are more prevalent in the midgut than in the hindgut. This pattern aligns with prior observations in the guts of *Dichelops melacanthus* and *Cephalotes rohweri* (Lanan et al., 2016; Prado & Zucchi, 2012). Although biofilm formation by gut symbionts has been documented across diverse animals, including fish, reptiles, insects, and humans (Agius et al., 2021; Burtseva et al., 2021; De Vos, 2015; Kim et al., 2014), biofilm-forming yeasts as bark beetle gut symbionts, and particularly in *I. sexdentatus* have yet to be reported. Furthermore, microscopic (SEM) visualisation of the dynamic interactions between gut yeast isolates and entomopathogenic fungi (EPF) or beetle-associated non-pathogenic ophiostomatoid fungi revealed distinct degradation and deformation of EPF mycelia. In contrast, no such structural disorientation was observed in the ophiostomatoid fungi. This suggests a distinct selective antagonism toward entomopathogenic fungi while maintaining a coexistence with wood-associated fungi in beetles. Interestingly, the yeast isolate *C. mississippiensis,* which exhibits chitinase and β-glucanase activities, showed extensive disruption of fungal cell walls and mycelial collapse (Toral et al., 2018). Moreover, the digestive CAZyme activities, such as amylase, cellulase, pectinase, and xylanase, of the yeast isolates suggest a contributory role in mycelial distortion and cell wall degradation. The antifungal and digestive enzyme activities of the studied yeasts isolated from various sources have been previously reported (Cheng et al., 2023; Daskaya-Dikmen et al., 2018; Kham et al., 2024; Lopes et al., 2018; Morais et al., 2020). Hence, the structural deformation of fungal mycelia could plausibly be due to the ability of the gut isolates to secrete extracellular antifungal and digestive enzymes. Nonetheless, biofilm formation in the midgut by the isolates may further potentiate both antifungal and digestive functions. Additionally, the non-entomopathogenic ophistomoid fungi that co-occur with bark beetles may be intrinsically less vulnerable under gut-like conditions, or effectively buffered by ecological compatibility (e.g., tolerance/neutrality within beetle–fungus consortia). This interpretation is consistent with reports that beetle-associated yeasts preferentially inhibit entomopathogenic fungi (Pineda-Mendoza et al., 2024). Therefore, the enzyme repertoire that disrupts EPF mycelia appears to spare non-entomopathogenic fungi, indicating strong selectivity. Such selective antagonism is likely adaptive, as it protects the beetles from pathogens while preserving non-pathogenic and potentially beneficial fungal associates. However, this hypothesis warrants further experimental validation to confirm the molecular underpinnings of this adaptive selectivity.

## Conclusion

This study demonstrates that *I. sexdentatus* and *I. acuminatus* maintain a stable core mycobiota throughout their life stages, which aids in nutrient acquisition, detoxification, and chemical signalling. However, mycobiome composition and diversity shift in response to beetle development, breeding environment, and interactions with trees. Beetle larvae displayed higher diversity than pupae and adults, with wild populations having richer fungal communities than lab-bred counterparts. The overlap between beetle and wood mycobiomes suggests a mutual microbial exchange during feeding, with saprotrophs dominating. Monoterpene blends were more detrimental to yeast symbionts than individual compounds, and selective antagonism was observed towards entomopathogenic fungi. Enzymatic assays further corroborate the rationale underlying such antagonistic activities. Nevertheless, these findings enhance our understanding of beetle-fungus interaction dynamics and highlight the need for integrated methods to explore their ecological roles. Future research should focus on unravelling these environmental interactions at the molecular level and exploring novel microbe-mediated bark beetle management strategies.

## Data Availability

The mycobiome dataset in the study is available under NCBI Bio-project PRJNA854812.

## Acknowledgements

We acknowledge the support of Dr. Roman Modlinger (FLD, CZU) during bark beetle collection from Czech forests. We thank Prof. Fredrik Schlyter (SLU, Sweden) for continuous encouragement and advices.

## Author Contributions

AK: Writing – original draft, Methodology, Visualization, Formal analysis, Data curation, Validation; SB: Writing– original draft, Methodology, Visualisation, Formal analysis; AC: Conceptualization, Writing – review & editing, Supervision, Resources, Investigation, Funding acquisition, Formal analysis, Data curation. JD: Methodology, Visualization, Writing – review & editing. JS: Methodology, Writing – review & editing; AR: Conceptualization, Writing – review & editing, Visualization, Supervision, Resources, Project administration, Investigation, Funding acquisition.

## Ethics statement

The manuscript presents research on animals that do not require ethical approval for their use in the study.

## Supplementary Figures

**Supplementary Figure 1** Rarefaction curves.

**Supplementary Figure 2** LefSe analysis-Host contribution. (A) Cladogram depicting significant fungal biomarkers in *I. acuminatus* wild beetles and respective wood samples (control wood and fed wood). (B) Cladogram illustrates significant fungal biomarkers among *I. sexdentatus* wild adult beetles and different wood types (control wood and fed wood). IAC- *I. acuminatus*; ISX- *I. sexdentatus*; WL- wild-collected; Ctrl. W- Control uninfested wood; Fed. W- Gallery wood.

**Supplementary Figure 3** LefSe analysis. Histogram revealing the LDA scores (threshold value set at log10>4) of significantly abundant fungal communities in various comparisons (A) Different developmental stages of *I. acuminatus* (larvae, pupae, adult), (B) Different life stages of *I. sexdentatus* (larvae, pupae, adult), (C) Lab-bred and wild adults of *I. acuminatus* and *I. sexdentatus*. (D) *I. acuminatus* wild adults compared with control wood and fed wood. (E) *I. sexdentatus* wild adults compared with control wood and fed wood. IAC- *I. acuminatus*; ISX- *I. sexdentatus*; WL- wild-collected; Ctrl. W- Control uninfested wood; Fed. W- Gallery wood.

**Supplementary Figure 4** Colonization of symbiotic yeast in the different gut compartments of *Ips sexdantatus*. (A) Colonization of symbiotic yeast in the midgut region. (B) Single yeast cells are localized in the midgut region. (C) Colonization of symbiotic yeast in the hindgut region. (D) Single yeast cells are localized in the hindgut region.

**Supplementary Figure 5** Biofilm-forming potential of symbiotic yeasts. (A) Biofilm formation by *Nakazawaea holstii*. (B) Biofilm formation by *Cyberlindera missipplensis*. (C) Biofilm formation by *Yamadazma mexicana*. (D) Biofilm formation by *Kuraishia molischiana*.

**Supplementary Figure 6** Individual morphology of symbiotic yeast. (A) Single-cell morphology of *Nakazawaea holstii.* (B) Single cell morphology of *Cyberlindera missipplensis*. (C) Single cell morphology of *Yamadazma mexicana.* (D) Single cell morphology of *Kuraishia molischiana*.

## Supplementary Tables

**Supplementary Table 1** Selected fungal primers used for the real-time quantitative PCR assay.

**Supplementary Table 2** List of fungal isolates.

**Supplementary Table 3** Composition of media used for different enzyme production.

**Supplementary Table 4** Alpha diversity indices. The data represent the mean value ± SE for five biological replicates across different life stages of two pine-feeding beetles and four biological replicates for the associated wood samples. (SE-Standard Error).

**Supplementary Table 5** ADONIS analysis using Bray-Curtis method to comprehend significant differences across developmental stages of both *Ips* pine beetles (ISX-*I. sexdentatus*, IAC-*I. acuminatus*) and wood samples (Df = degree of freedom, SS = sums of squares of deviations, MS = SS/Df, F. Model = F-test value, R2 = the ratio of grouping variance and total variance).

**Supplementary Table 6** ANOSIM analysis revealing significant variation across different developmental stages of both *Ips* pine beetles (ISX-*I. sexdentatus*, IAC-*I. acuminatus*). The R values closer to 1.0 denote significant differences in fungal communities between different developmental stages of both beetles. P value < 0.05 denotes a statistically significant difference.

## Supplementary Excels

**Supplementary Excel 1** Raw and clean reads.

**Supplementary Excel 2** ASV table representing the relative abundance of fungal community in two Ips beetles *I. sexdentatus* (ISX) and *I. acuminatus* (IAC), along with their respective wood samples.

**Supplementary Excel 3** Relative abundance of top 10 fungal order.

**Supplementary Excel 4** Core and Unique mycobiome in *I. acuminatus* (IAC)

**Supplementary Excel 5** Core and Unique mycobiome in *I. sexdentatus* (ISX)

**Supplementary Excel 6** Common and unique mycobiome among different stages of *I. sexdentatus* (ISX), *I. acuminatus* (IAC).

**Supplementary excel 7** Core and unique fungal communities illustrating the effect of lab breeding on the pine-feeding beetle mycobiome

**Supplementary excel 8** Common and unique mycobiome representing the host contribution to the beetle mycobiome.

**Supplementary Excel 9** Alpha diversity indices.

**Supplementary Excel 10** q-PCR statistics

